# A single-sample workflow for joint metabolomic and proteomic analysis of clinical specimens

**DOI:** 10.1101/2023.11.07.561857

**Authors:** Hagen M. Gegner, Thomas Naake, Karim Aljakouch, Aurelien Dugourd, Georg Kliewer, Torsten Müller, Dustin Schilling, Marc A. Schneider, Nina Kunze-Rohrbach, Thomas G.P. Grünewald, Rüdiger Hell, Julio Saez-Rodriguez, Wolfgang Huber, Gernot Poschet, Jeroen Krijgsveld

## Abstract

Understanding the interplay of the proteome and the metabolome aids in understanding cellular phenotypes. To enable more robust inferences from such multi-omics analyses, combining proteomic and metabolomic datasets from the same sample provides major benefits by reducing technical variation between extracts during the pre-analytical phase, decreasing sample variation due to varying cellular content between aliquots, and limiting the required sample amount. We evaluated the advantages, practicality and feasibility of a single-sample workflow for combined proteome and metabolome analysis. In the workflow, termed MTBE-SP3, we combined a fully automated protein lysis and extraction protocol (autoSP3) with a semi-automated biphasic 75% EtOH/MTBE extraction for quantification of polar/non-polar metabolites. Additionally, we compared the resulting proteome of various biological matrices (FFPE tissue, fresh-frozen tissue, plasma, serum and cells) between autoSP3 and MTBE-SP3. Our analysis revealed that the single-sample workflow provided similar results to those obtained from autoSP3 alone, with an 85-98% overlap of proteins detected across the different biological matrices. Additionally, it provides distinct advantages by decreasing (tissue) heterogeneity by retrieving metabolomics and proteomic data from the identical biological material, and limiting the total amount of required material. Lastly, we applied MTBE-SP3 to a lung adenocarcinoma cohort of 10 patients. Integrating the metabolic and proteomic alterations between tumour and non-tumour adjacent tissue yielded consistent data independent of the method used. This revealed mitochondrial dysfunction in tumor tissue through deregulation of OGDH, SDH family enzymes and PKM. In summary, MTBE-SP3 enables the facile and confident parallel measurement of proteins and metabolites obtained from the same sample. This workflow is particularly applicable for studies with limited sample availability and offers the potential to enhance the integration of metabolomic and proteomic datasets.

## Introduction

Since proteins and metabolites constitute a rich representation of the cell’s phenotype, their collective analysis has contributed to elucidate cellular mechanisms in multiple scenarios. In a clinical setting, integrating proteomic and metabolomic data with genomic and transcriptomic profiles has the potential to significantly enhance personalised medicine strategies and to diagnose and stratify patients (Kowalczyk *et al*, 2020). Integrative strategies approaches that combine various omics approaches further enhance the capability to study the interplay between regulatory layers and provide insights into complex and multifactorial pathologies, such as cancer (Yoo *et al*, 2018).

Metabolomic and proteomic sample preparation workflows have traditionally focused on optimising extraction conditions to maximise metabolite or protein coverage (Hughes *et al*, 2014; Cai *et al*, 2022; Varnavides *et al*, 2022; Gegner *et al*, 2022a). More recently, this has been extended with efforts to standardise these methodologies, ideally in an automated fashion, driven by the need to minimise inconsistencies introduced by sample handling, especially in large sample cohorts (Müller *et al*, 2020; Leutert *et al*, 2019). For metabolomics, biphasic extractions, utilising either ethanol or methanol combined with methyl-tert-butylether (MTBE), showed advantages over using chloroform or monophasic extractions by exhibiting higher coverage, increased extracted metabolite concentration and robustness (Erben *et al*, 2021; Gegner *et al*, 2022a). Similarly to metabolomics, the proteomic workflow using single-pot solid-phase-enhanced sample preparation (SP3) on a liquid handling robot for automated processing (autoSP3) revealed several advantages over various existing automated proteomics workflows (Mueller et al., 2020). These include the ability to process low-input samples, to increase the reproducibility of the proteomic output and to reduce the variability in protein quantification in a cost-effective manner.

In conventional approaches for combined proteomic and metabolomic studies, samples are often prepared separately from different specimens. This is not an ideal approach as any inconsistencies between the proteomic and metabolomic data may be incorrectly interpreted as regulatory interactions between these two layers, while in fact this might arise from sample variability. For instance, differences in pre-analytical sample handling (e.g. time and temperature of storage) (Gegner *et al*, 2022b) may be likely to occur if proteomic and metabolomic sample preparation is conducted in different labs. Therefore, consistency between proteomic and metabolomic data may be significantly enhanced if they are generated from physically the same sample, thus benefiting clinical or mechanistic interpretation of the combined data (Garikapati *et al*, 2022; Bayne *et al*, 2021; Zougman *et al*, 2020; Nakayasu *et al*, 2016). In addition, utilising single-sample workflows also offers several other advantages, such as minimising the pre-analytical variability and reducing sample heterogeneity related to factors such as tumour content. Furthermore, the required total sample amount can be limited.

These benefits have prompted several studies to develop single-sample workflows for combined proteomic, metabolomic and in some cases lipidomic analysis (Zougman *et al*, 2020; Nakayasu *et al*, 2016; Coman *et al*, 2016). Yet, with few exceptions (Garikapati *et al*, 2022) these studies focused on the analysis of one sample type (e.g. cells, tissue or plasma), and thereby the universal applicability to all biological matrices remains unclear. In addition, these approaches largely employ manual sample handling procedures, although it has been noted that several steps are amenable for automation (e.g. cell lysis, protein digestion) to enhance reproducibility (Gutierrez *et al*, 2018).

SP3 has become a broadly used method for proteomic sample preparation because of its wide applicability, high sensitivity, ease of use, and low cost (Hughes et al, 2014; Varnavides *et al*, 2022; Sielaff *et al*, 2017), that we previously implemented on a robotic platform as autoSP3 (Müller *et al*, 2020). Here we aimed to assess the performance of a one-sample strategy that combines autoSP3 with an optimised approach for metabolomics (Gegner *et al*, 2022a). In addition, we aimed to apply the combined workflow to different biological matrices, and to benefit from the capacity of SP3 for automated proteomic sample preparation to enhance standardisation of proteo-metabolomic studies. In particular, we subjected several sample types (formalin-fixed paraffin-embedded (FFPE) tissue, fresh-frozen tissue, plasma, serum, and cells) to bi-phasic extraction of metabolites with MTBE (Gegner *et al*, 2022a), resulting in a precipitated protein pellet that was subsequently used as a direct input for the proteomic workflow utilising automated and parallelized sonication and protein clean-up by autoSP3.

We demonstrate that the proteomic data generated by the MTBE-SP3 approach is highly consistent with the original autoSP3 method. Further extending its utility, the MTBE-SP3 approach offers a universal applicability across a broad range of biological matrices. Next, we applied the combined workflow on a lung adenocarcinoma patient cohort and used a novel network approach to determine that consistent metabolic and proteomic alterations were observed between tumour and non-tumour adjacent tissue, independent of the method that was used for proteomics (autoSP3 or MTBE-SP3). Hence, MTBE-SP3 is a powerful and robust method for integrated metabolomic and proteomic studies performed on the same sample that can be employed for universal applications in diverse biological matrices.

## Results

### The single-sample workflow yields similar results compared to autoSP3

Here, we aimed to establish a strategy that combines two methods that had been individually optimised for proteome and metabolome analysis, i.e. SP3 and EtOH/MTBE, respectively, for integrated proteo-metabolomic analysis of physically the same sample. In particular, we used an organic solvents-based extraction to release metabolites, leaving a protein-containing residue that we used as an input for SP3. In more detail, we applied a bi-phasic extraction with MTBE and 75% ethanol (EtOH) that precipitates proteins as a pellet and generates an upper organic phase containing lipids, and a lower aqueous phase containing polar metabolites (Figure 1A). The liquid extract, containing the upper and middle phase (Figure 1A), was transferred to a new reaction tube, dried, and resuspended for downstream targeted metabolomics via the Biocrates MxP Quant 500 kit (Gegner *et al*, 2022a) while the pellet, containing the precipitated proteins (Figure 1A), was used as direct input for the standard autoSP3 workflow (Müller *et al*, 2020), followed by a DDA approach on a timsTOF Pro mass spectrometry for proteome analysis.

**Figure 1:**
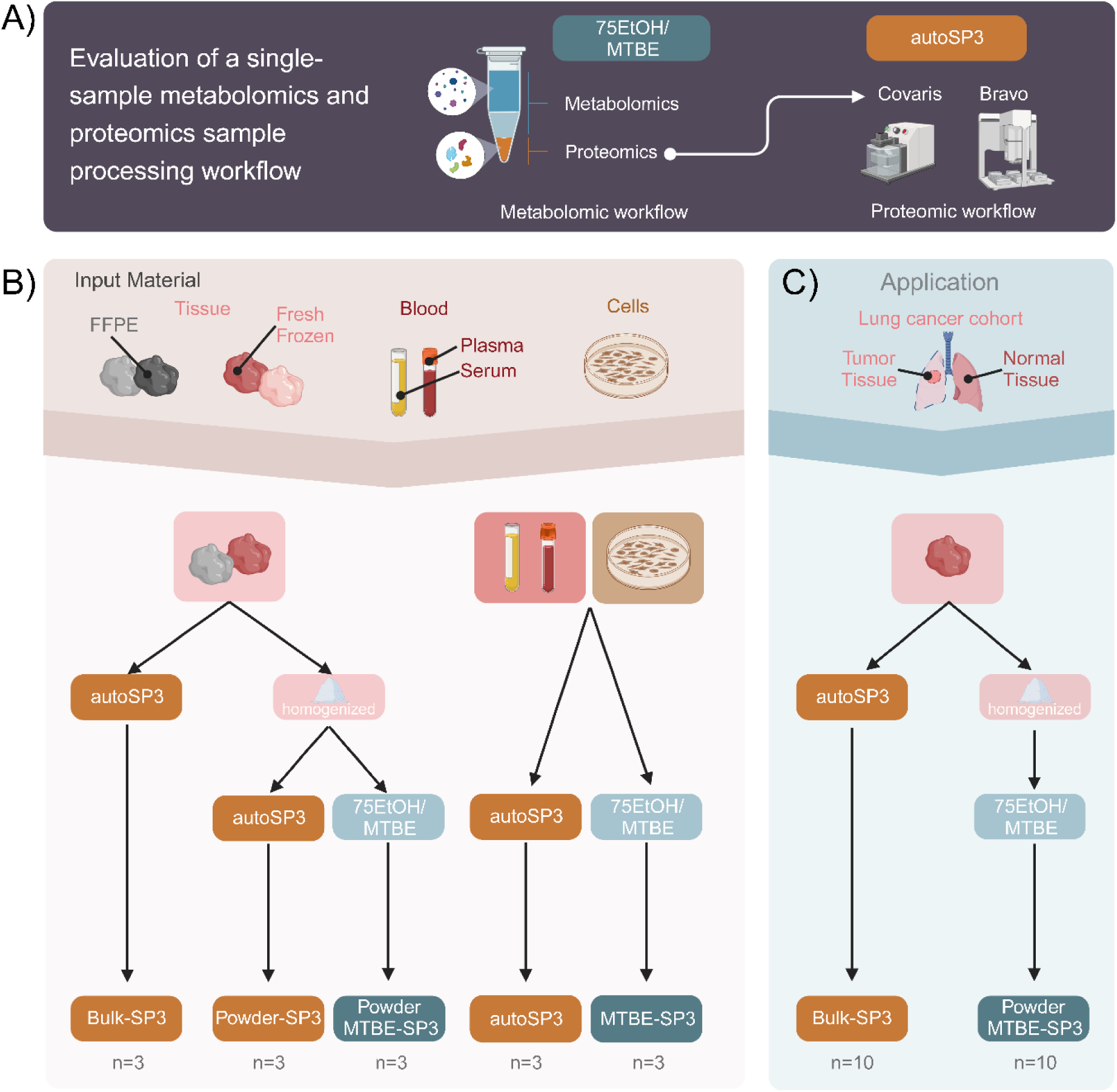
Overview of experimental setup. A) Proteins were extracted using two different methods: the established autoSP3 method (Müller *et al*, 2020) and the single-sample workflow via 75EtOH/MTBE extraction followed by autoSP3 (MTBE-SP3). B) The two extraction methods were tested and compared for several biological matrices (FFPE tissue, fresh-frozen tissue, cells, plasma, and serum). For FFPE and fresh-frozen tissue samples, tissue pieces (bulk) were either used as a direct input for autoSP3 or were cryo-pulverised and homogenised (powder). The powder was then used either as a direct input for autoSP3 (Powder-SP3) or subjected to the 75EtOH/MTBE extraction followed by autoSP3 (Powder MTBE-SP3). For serum, plasma and cells, samples were used either as direct input for autoSP3 or the biphasic 75EtOH/MTBE extraction followed by autoSP3 (MTBE-SP3). C) To test the concordance between biological interpretations, both extraction methods were tested on a lung adenocarcinoma cohort and the resulting proteomes were compared.

This single-sample extraction method (MTBE-SP3) was tested on five different biological matrices: FFPE tissue, fresh-frozen tissue, plasma, serum, and cells (see methods for sample origin and further details on the biological matrices). Crucially, we compared the MTBE-SP3 extraction approach to the original autoSP3 method, which extracts proteins using an SDS-containing buffer and does not include a protein precipitation step (Müller *et al*, 2020) to assess completeness and potential bias in proteome coverage. Here, we consider technical replicates as repeat applications of the same extraction method: for each biological matrix we acquired three samples per extraction method (autoSP3, MTBE-SP3; Figure 1) that were analysed for proteomics. For FFPE and fresh-frozen tissues, proteins were extracted from bulk as a direct input to autoSP3 (Bulk-SP3) or from cryo-pulverised and homogenised tissue (Powder-SP3) and following the 75EtOH/MTBE extraction step (Powder-MTBE-SP3). Bulk FFPE and fresh-frozen samples were physically distinct tissue pieces, while samples from homogenised samples were taken from the same homogenate. For plasma, serum, and cell samples, proteins were extracted from the bulk (autoSP3) or following the 75EtOH/MTBE extraction step (MTBE-SP3).

In a first analysis, we assessed the recovery of proteins based on the MaxQuant identification to check whether autoSP3 and MTBE-SP3 methods obtain similar sets of proteins. In terms of recovery of proteins in the two extraction methods, both protocols showed a high overlap of detected proteins (Figure 2B, see also Figure 2C for fresh-frozen tissue and Supplementary Figure S1 for data in other sample types). Looking at the shared protein identifiers after MaxQuant identification, the MTBE-SP3 method showed high overlap of detected proteins compared to the Powder-autoSP3/Bulk-autoSP3 in the FFPE (85%) and fresh-frozen samples (89.4%) and high overlap compared to autoSP3 in cells (97.6%), serum (90%) and plasma (91%). This indicates very similar efficiency of the extraction methods, which was also confirmed by the highly comparable LFQ intensity range in the respective proteomic datasets (Figure 2A).

**Figure 2:**
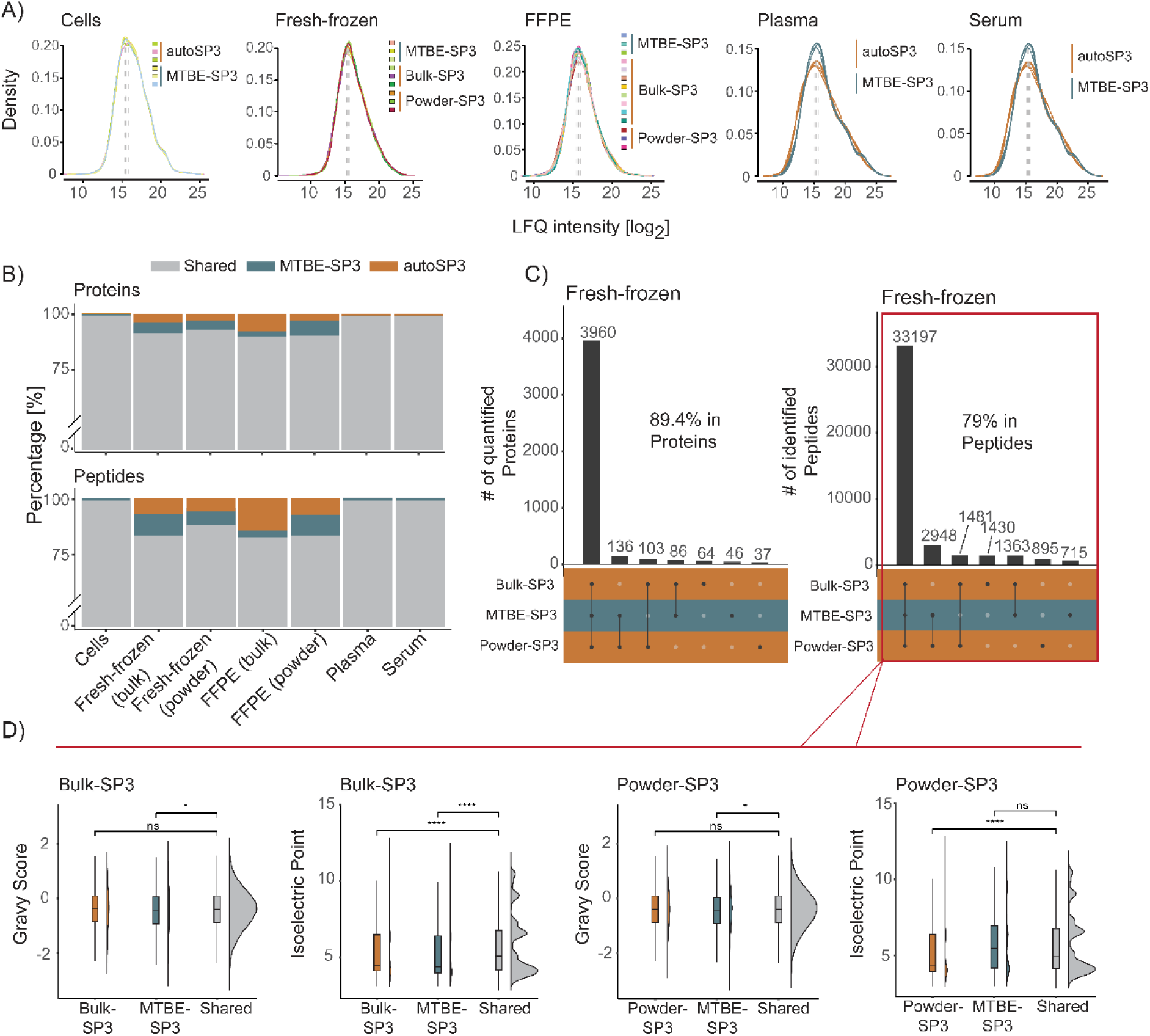
Intensities and overlap across all sample types. A) Densities of log-transformed LFQ intensities for the replicates in all sample types. B) Bar chart illustrating the percentage of shared (common) and unique quantified proteins and peptides in MTBE-SP3 and autoSP3. C) Joint and disjoint proteins and peptide sets in fresh-frozen samples. While some of the proteins and peptides were uniquely detected in one of the extraction methods (MTBE-SP3, autoSP3), the majority of proteins and peptides were detected in both methods. The numbers (in %) indicate the proportion of the largest set relative to the total number of proteins and peptides. D) GRAVY and isoelectric point scores for proteins for the sets autoSP3/MTBE-SP3.

Next, we evaluated whether MTBE-SP3 yields concordant results to the established autoSP3 protocol by the following measures: *i*) the number of differentially expressed proteins between the two extraction methods, *ii*) the correlation of log-transformed intensities of technical replicates, *iii*) and the precision of measurements expressed by the coefficient of variation (CV) of the technical replicates. *i*) We found a variable, but generally low number of proteins that differed in abundance (fresh-frozen tissue, powder: 0%; FFPE tissue, powder: 0%; cells: 1.1%; serum: 4.6%; plasma: 14.4%; FFPE tissue, bulk: 15.1%; fresh-frozen tissue, bulk: 19.3%). Especially the homogenised tissues showed no abundance differences between the two extraction methods, indicating their equivalent performance. In contrast, these numbers were higher for bulk samples, indicating that, as expected, non-homogenized samples exhibit higher variability in their protein content (Table 1). *ii*) For FFPE and fresh-frozen samples, the correlation analysis between technical replicates revealed high CVs between MTBE-SP3 and (homogenised) auto-SP3 (average R^2^=0.80, SD=0.05 for FFPE, and R^2^=0.91, SD=0.02 for fresh frozen), and to a lesser extent between MTBE-SP3 and Bulk-SP3 (average R^2^= 0.73, SD=0.06 for FFPE and R^2^=0.82, SD=0.04 for fresh frozen). For plasma, serum and cells high coefficients were obtained between MTBE-SP3 and auto-SP3 with an average R^2^=0.89, SD=0.03 for plasma, R^2^=0.92, SD=0.01 for serum, and R^2^=0.92, SD=0.01 for cells. (Supplementary Figure S2). *iii*) Similarly to autoSP3, MTBE-SP3 showed low CVs for liquid (plasma, serum), pulverised (fresh-frozen and FFPE tissue), and other matrices (cells, bulk fresh-frozen, and bulk FFPE tissue). While the differences in CV were significantly different between MTBE-SP3 and autoSP3 for most of the sample types (except serum, α < 0.05, no FDR correction), the effect size was generally low in absolute terms (Table 1).

**Table 1:**
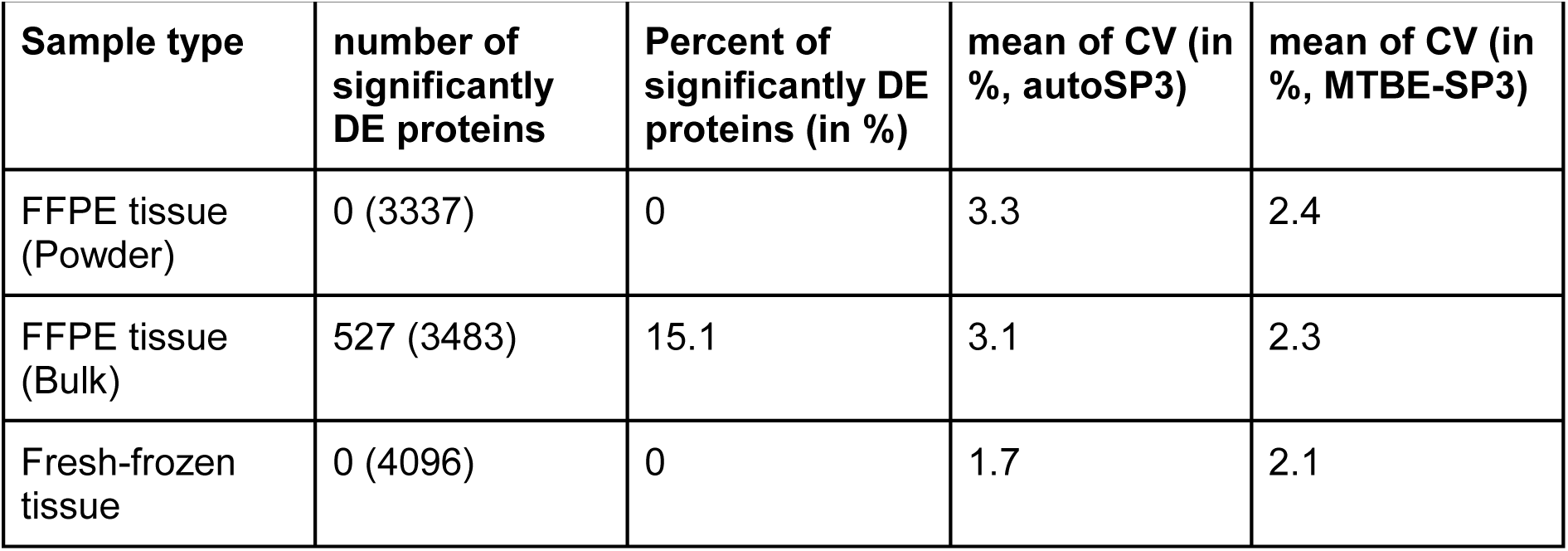
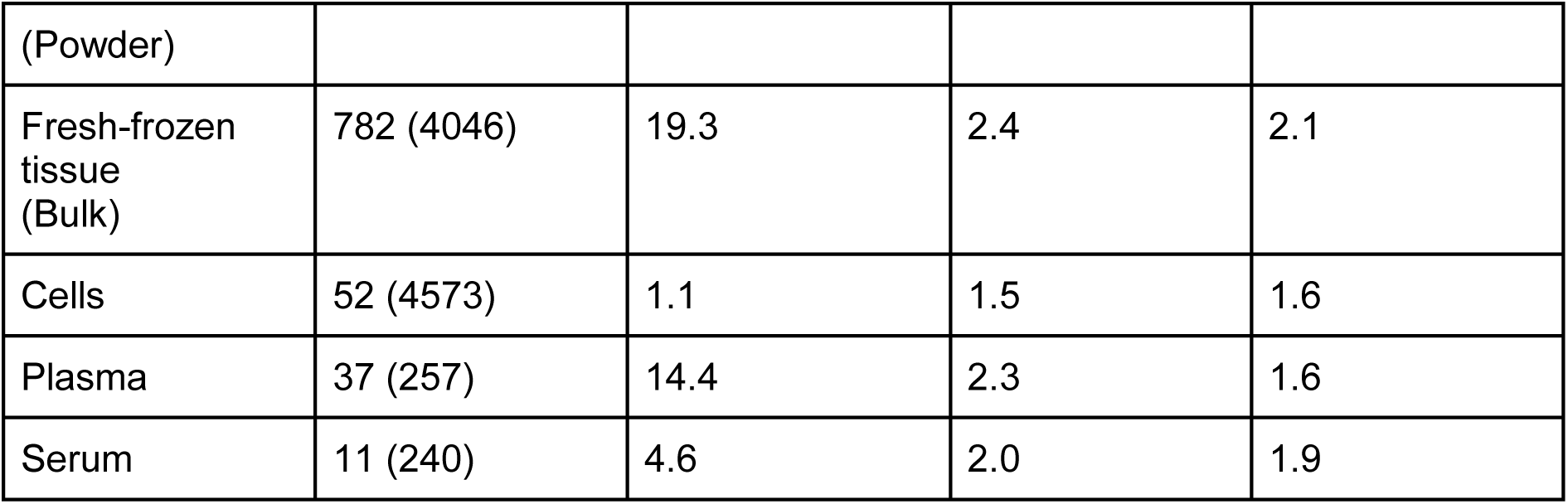
Differentially expressed proteins between autoSP3 and MTBE-SP3 and coefficient of variation (CV) between replicates. For all tissues, DE proteins were determined using linear models by testing differences between the replicates extracted with autoSP3 *vs.* the replicates extracted by MTBE-SP3. Reported here are the number of significantly DE proteins (α < 0.05 after FDR correction) for each experiment. The number in brackets shows the total number of tested proteins. The percent of significantly DE proteins was calculated from the number of significantly DE proteins and total number of tested proteins. The CV values were calculated from the mean of and standard deviation between technical replicates of each condition, e.g. of the autoSP3-derived technical replicates and the MTBE-SP3-derived technical replicates of the cell dataset. CV values are reported in percent. CV: coefficient of variation; DE: differentially expressed.

Moreover, we devised an R package (PhysicoChemicalPropertiesProtein, available via www.github.com/tnaake/PhysicoChemicalPropertiesProtein) to calculate two important parameters, the isoelectric point and GRAVY (grand average of hydropathy) scores, to scrutinise potential differences in extraction efficiencies regarding physico-chemical properties (Figure 2D, Supplementary Figure S3). To that end, we correlated the values of the GRAVY/isoelectric point scores for proteins with the t-values from differential expression analysis. The t-values were regarded as a measure of how differently abundant proteins are for a given extraction method. The homogenous samples (FFPE (powder), cells, plasma, and serum), showed no clear association between the GRAVY/isoelectric point scores and t-values (Spearman ρ correlation coefficients close to 0). These small correlation coefficients were not statistically significantly different from 0, indicating that there is no bias in physico-chemical properties of proteins in the tissues FFPE (powder), cells, plasma, and serum. FFPE (bulk) and fresh-frozen tissue (powder and bulk) showed a moderate positive correlation between GRAVY scores and t-values (Table 2). This suggests that more hydrophobic proteins were detected in higher abundance in these matrices in autoSP3 compared to the MTBE-SP3 extraction. Accordingly, GO terms related to the membrane system were differentially expressed between autoSP3 and MTBE-SP3 extraction in fresh-frozen tissue (bulk), while FFPE (bulk) showed enrichment of terms related to the cytoskeleton and DNA/RNA-related processes (Supplementary Figure S4). These differences may be explained from the fact that, by necessity, bulk samples were prepared from disparate tissue pieces which may have differed in composition. Therefore, in conclusion, our data show that depending on the tissue type MTBE-SP3 is equivalent to autoSP3 with regard to the proteome coverage that is obtained across a variety of sample types, with no noticeable (e.g. for fresh-frozen tissue, powder; FFPE tissue, powder; or cells) or moderate selectivity (e.g. FFPE tissue, bulk, fresh-frozen tissue, bulk) in protein extraction.

**Table 2:**
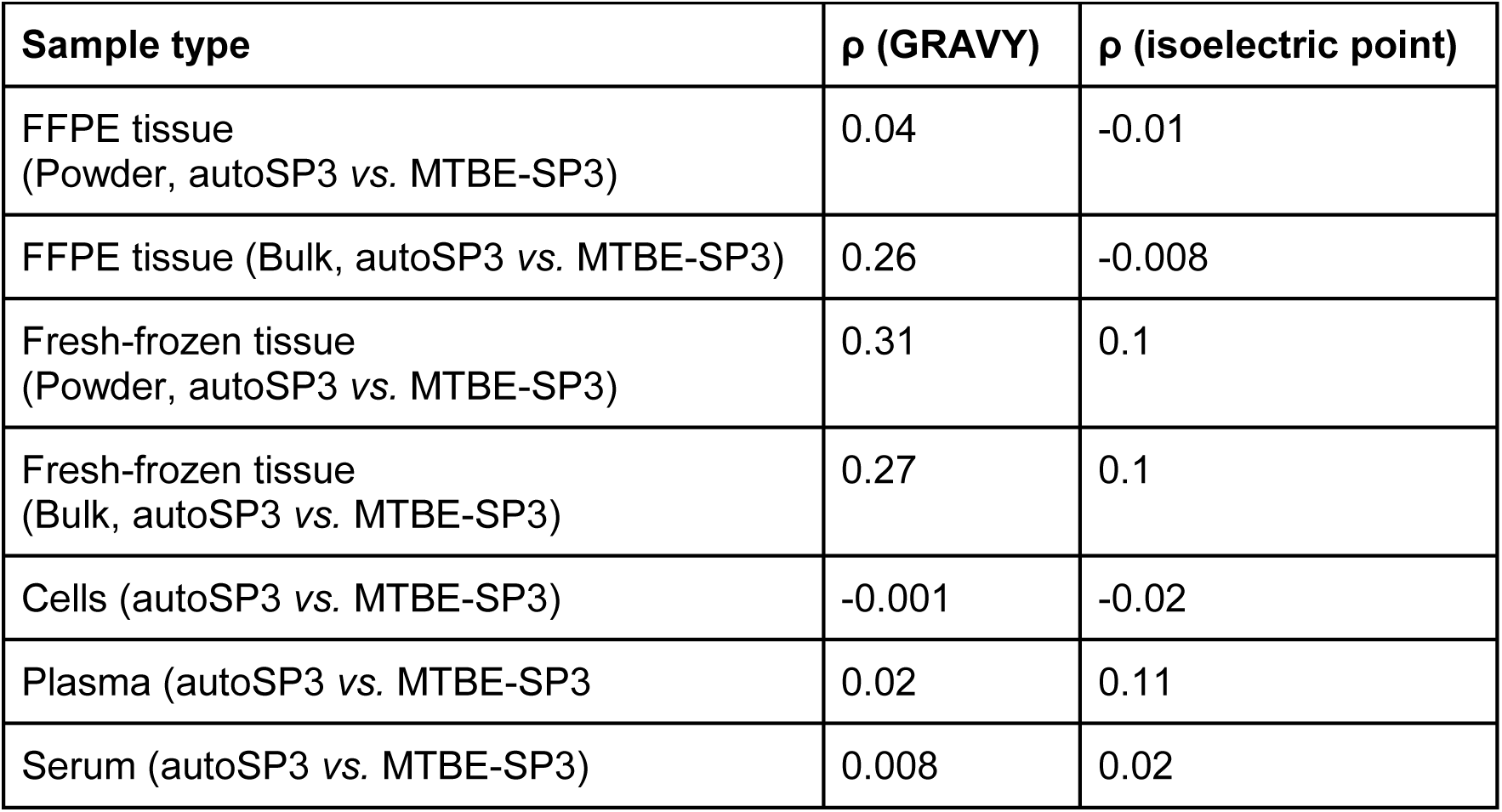
Spearman ρ correlation coefficients between GRAVY scores or isoelectric point values and t-values. GRAVY scores and isoelectric point values were derived from the amino acid sequences of proteins. For each tissue, the t-values from differential expression analysis derived from the protein abundances were correlated using Spearman’s Rank correlation against the GRAVY scores or isoelectric point values.

### Applying MTBE-SP3 on a lung adenocarcinoma cohort yields similar results compared to autoSP3

To demonstrate the advantages of the MTBE-SP3 workflow, we applied it in a combined proteome and metabolome analysis in a lung adenocarcinoma cohort. The cohort consisted of fresh-frozen samples from ten patients of paired tumorous tissue (TT) and non-tumorous adjacent tissue (NAT). A particular aim was to assess if similar biological conclusions can be reached in the comparison of these tissue regions when using autoSP3 or MTBE-SP3 for proteome analysis, despite minor differences that may exist between these methods. In addition, using MTBE-SP3, we performed broad-scale targeted metabolomics via MxP Quant 500 (Biocrates). In total, across all samples we quantified 6326 proteins in a single-shot DDA approach using a timsTOF Pro mass spectrometer. After filtering the data, proteomic data was available for 3010 protein features with quantitative information in >50% of the samples, which were included for further analysis. The metabolomic dataset contained concentrations for 405 metabolites after applying the filtering steps based on the MetIDQ-derived quality scores (see Materials & Methods for further details).

To address if autoSP3 and MTBE-SP3 yield similar quantification results we determined if protein abundances differ when using them for protein extraction from either NAT or TT samples. Analysis of 10 *vs.* 10 NAT tissue pieces processed by autoSP3 and MTBE-SP3, respectively, identified 3010 proteins of which 809 showed a difference in abundance (α < 0.05 after FDR correction). For TT samples, 553 out of 3010 proteins showed an abundance difference. To test whether this difference may be explained by tissue heterogeneity, we run linear models for the two extraction methods separately on random, equally split partitions of samples. This analysis did not show any differentially expressed proteins for either autoSP3 or MTBE-SP3 (α < 0.05 after FDR correction), indicating that tissue heterogeneity is not governing the observed differences. This suggests that slight differences exist between both methods for this type of samples, although fold changes were mostly modest. This is not necessarily problematic as long as no bias is introduced that skews biological differences between samples that are analysed with either method. To test this, we assessed if autoSP3 and MTBE-SP3 yield the same sets of differentially expressed proteins between NAT and TT samples. When looking at the NAT *vs.* TT differences adjusting for the autoSP3 and MTBE-SP3 methods (i.e., considering the differences between NAT_autoSP3_ *vs.* TT_autoSP3_ and NAT_MTBE-SP3_ *vs.* TT_MTBE-SP3_), only two proteins were significantly different (PDLIM2 and PRPF40A, α < 0.05 after FDR correction, Figure 3A), indicating the equivalence of both sample preparation methods.

**Figure 3:**
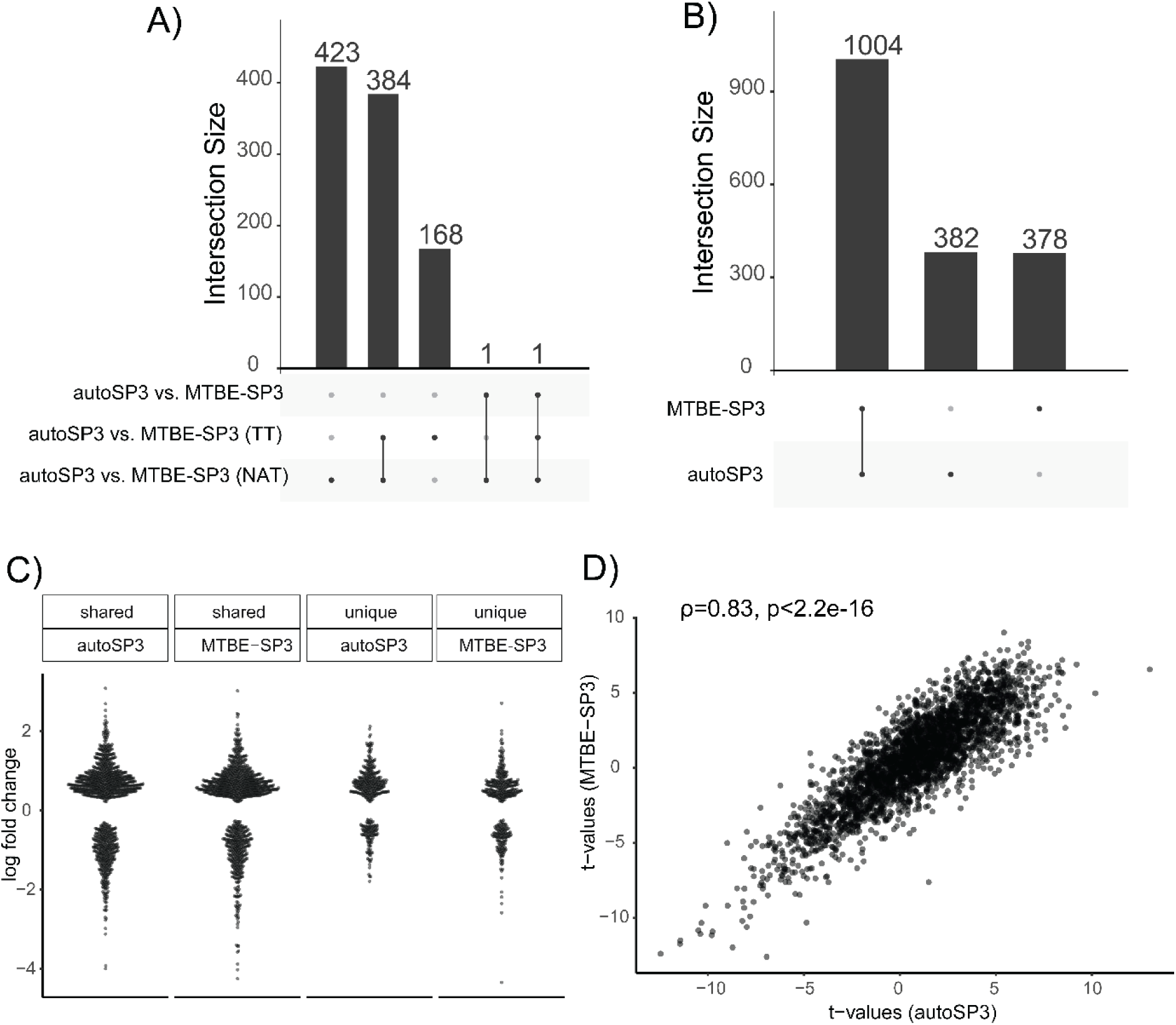
Differential expression analysis for lung adenocarcinoma cohort (proteomics). A) UpSet plot of significant protein features for contrast autoSP3 *vs.* MTBE-SP3 (α < 0.05 after FDR correction). The DE analysis was performed on the sets corresponding to autoSP3 *vs.* MTBE-SP3 for NAT samples, autoSP3 *vs.* MTBE-SP3 for TT samples, and autoSP3 *vs.* MTBE-SP3 for the entire sample set. B) UpSet plot for contrast TT *vs.* NAT. The DE analysis was performed on the sets derived from autoSP3 and MTBE-SP3 extraction. C) Beeswarm plot of log fold changes. The sets correspond to the protein sets from panel B: ‘shared autoSP3’ corresponds to the log fold changes of the 1004 proteins in the autoSP3 dataset, ‘shared MTBE-SP3’ to the log fold changes of the 1004 proteins in the MTBE-SP3 dataset, ‘unique autoSP3’ corresponds to the log fold changes of the 382 proteins in the autoSP3 dataset, and ‘unique MTBE-SP3’ corresponds to the log fold changes of the 378 proteins in the MTBE-SP3 dataset. The absolute log fold changes in the shared sets are higher compared to the unique sets (autoSP3: W = 239420, p-value < 4.2e-13; MTBE-SP3: W = 230510, p-value < 3.6e-10; Wilcoxon rank sum test with continuity correction, no adjustment for multiple testing). D) Scatter plot between t-values from MTBE-SP3 and t-values from autoSP3. The Spearman’s rank correlation ρ between the two sets of t-values is 0.83 (p-value < 2.2e-16, no FDR correction). DE: differential expression/differentially expressed. NAT: non-tumorous adjacent tissue. TT: tumorous tissue.

We next determined the overlap among the proteins that were differentially expressed between NAT *vs.* TT, as obtained by autoSP3 and MBTE-SP3. The extraction methods detected 1386 (autoSP3) and 1382 proteins (MTBE-SP3) to be differentially expressed between NAT and TT (α < 0.05 after FDR correction). Of these, 1004 proteins were shared among autoSP3 and MTBE-SP3, while 382 (autoSP3) and 378 (MTBE-SP3) were uniquely differentially expressed in each method (Figure 3B). The considerably lower number of statistically differentially expressed proteins above (NAT *vs.* TT adjusting for the autoSP3 and MTBE-SP3 methods) compared to the high number of unique proteins for each method tested individually can be explained by the further introduction of variation and higher number of levels of fitted cofactors when adjusting for the two extraction methods. The magnitude of the fold-change among the 1004 shared proteins was higher compared to the 382 and 378 proteins that were unique to autoSP3 and MTBE-SP3, respectively (autoSP3: Wilcoxon’s W = 239420, p-value < 4.2e-13; MTBE-SP3: Wilcoxon’s W = 230510, p-value < 3.6e-10; Wilcoxon rank sum test with continuity correction, no adjustment for multiple testing, Figure 3C), indicating that main differences were captured by both methods. The t-values of the contrast NAT *vs.* TT for autoSP3 and MTBE-SP3 showed a high correlation (Figure 3D, ρ = 0.83, p-value < 2.2e-16, no FDR correction) indicating that both autoSP3 and MTBE-SP3 detected the same differential expression patterns between NAT *vs.* TT. Thus, although autoSP3 and MTBE-SP3 show slight differences in sampling proteomes from these tissues, they yield similar results when comparing differences between samples (here NAT *vs.* TT) adjusting for the extraction method. Taken together, the results indicate that autoSP3 and MTBE-SP3 perform similarly in quantifying proteome differences in complex clinical tissues.

### Integration of metabolomic and proteomic data

For the ten patients of the lung adenocarcinoma cohort, we additionally acquired metabolomic information using the Biocrates MxP Quant 500 kit. After performing quality control, the dataset contained information on the levels of 405 metabolites in the NAT and TT samples. Subsequently, we analysed the metabolomics dataset in conjunction with the MTBE-SP3 proteomics dataset, acquired from physically the same aliquot of the samples, and the autoSP3 proteomics dataset, acquired from a different aliquot of the samples (Figure 1C). To characterise the coherence of the proteomic and metabolomic data at the level of biological processes, we determined if MSigDB hallmark enrichment scores computed from proteomic and metabolomic data were correlated and checked if this correlation differed when proteomic data were obtained by MTBE-SP3 or autoSP3. This showed notably that the hallmark scores were highly correlated (0.83 to 0.94 Pearson’s R) when considering only proteins, and that the inclusion of metabolites did not affect the hallmark scores much (Figure 4A). Indeed, the number of measured metabolite features that could be mapped to metabolic pathways was not large enough to affect the correlation based on proteins. Nonetheless, we compared the hallmark scores that could be obtained specifically from proteomic or metabolic data, showing an average Pearson correlation of only 0.2 and 0.15 for MTBE-SP3 and autoSP3 proteomic data, respectively (Figure 4B). This low correlation is consistent with the notion that metabolic abundance usually correlates poorly with the abundance of metabolic enzymes, even in the same pathways, further supporting that metabolomic data allows to generate complementary insights in combination with proteomic data. Furthermore, we observed no significant difference between the correlation coefficients of the MTBE-SP3 and the autoSP3 datasets (Figure 4B, Student t-test p-value = 0.53, df = 3), indicating that both datasets are similar.

**Figure 4:**
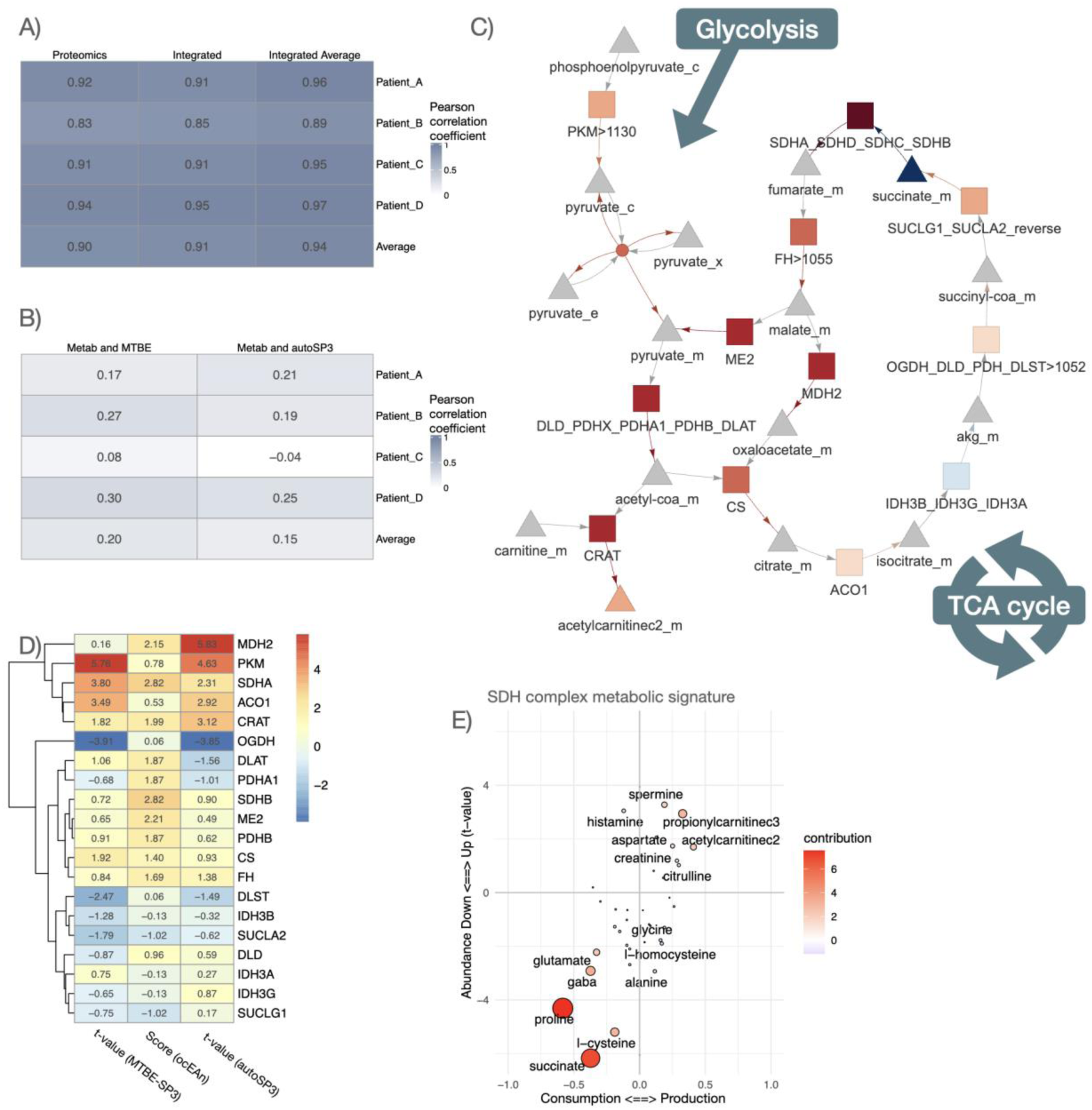
Comparison of proteomic and metabolomic integration between MTBE-SP3 and autoSP3. A) Pearson correlation coefficients between MTBE-SP3 and autoSP3 (*i*) proteomic MSigDB hallmark enrichment scores, (*ii*) integrated proteomic+metabolomic MSigDB hallmark enrichment scores, and (*iii*) averaged proteomic and metabolomic MSigDB hallmark enrichment scores. Hallmark enrichment scores were calculated using the decoupler package and represent the number of standard deviations away from the mean of an empirical null distribution of scores for a given hallmark. The colour gradient represents the correlation coefficient. B) Pearson correlation coefficients between MTBE-SP3 proteomic and metabolomic MSigDB hallmark enrichment scores (left column), and Pearson correlation coefficients between autoSP3 proteomic and metabolomic MSigDB hallmark enrichment scores (right column). Hallmark enrichment scores were calculated using the decoupler package and represent the number of standard deviations away from the mean of an empirical null distribution of scores for a given hallmark. C) Representation of the TCA cycle main enzymes and metabolites in ocEAn. Arrows represent consumptions (reactant to enzyme) and productions (enzymes to product) of metabolites. Colours represent positive (red, over-production and consumption) and negative (blue, under-production and consumption) metabolic ocEAn signature imbalance (signatures are defined as the sets of metabolites that are found upstream and downstream of a given enzyme in the whole metabolic reaction network). D) Heatmap displaying the t-values of TCA enzyme abundance changes between lung TT and NAT for the autoSP3 and MTBE-SP3 dataset, and ocEAn metabolic imbalance scores estimated from the differential metabolomic abundances between lung tumour and healthy tissue. E) Scatter plots representing the differential metabolomic abundances upstream (consumption) and downstream (production) of the SDH enzyme complex. The x-axis represents the ocEAn score, while the y-axis represents the corresponding t-value for a given enzyme (TT *vs.* NAT).

We then looked for more specific connections between enzymes and the overall metabolic deregulation profiles of tumours, and we assessed if they differ between MTBE-SP3 and autoSP3 datasets. The ocEAn package allows to explore connections between metabolites and metabolic enzymes beyond their direct interactions: ocEAn provides weighted interactions for all possible metabolites and enzymes of a reduced functional genome-scale metabolic network, where weights represent relative distances between metabolites and enzymes in the reaction network (Sciacovelli *et al*, 2022). ocEAn was used to systematically explore metabolites upstream and downstream of metabolic enzymes, in order to determine which of those showed the most imbalanced metabolic abundance signatures between TT and NAT samples, i.e. enzymes that show very different metabolic abundance profile changes upstream and downstream of their respective reactions (Figure 4C). Such imbalance can help to pinpoint metabolic bottlenecks in the metabolic reaction network, which can be more easily interpreted functionally than single metabolite abundance changes can. This notably showed that the succinate dehydrogenase (SDH) metabolic enzyme complex (composed of SDHA, SDHB, SDHC and SDHD), which converts succinate to fumarate as part of the Krebs cycle, was the most significantly imbalanced metabolic reaction according to metabolic deregulation in TT samples (Figure 4D, Figure 4E). Indeed, Figure 4E shows that the abundance of proline and succinate, which are consumed upstream of the SDH complex, are also significantly down-regulated (thus located in the lower left quadrant), while the abundance of spermine, propionylcarnitine and acetylcarnitine, which are produced downstream of the SDH complex, is significantly increased (thus located in the upper right quadrant). Interestingly, the MTBE-SP3 and autoSP3 datasets showed a significant up-regulation of the SDHA complex subunit in TT, albeit more significant in the MTBE-SP3 dataset (MTBE-SP3: t-value = 3.80, p-value = 0.001 after FDR correction; autoSP3: t-value = 2.31, p-value = 0.04 after FDR correction). The marginal accumulation of carnitine conjugates, such as propionyl-carnitine and acetyl-carnitine (p-value = 0.06 and 0.27 respectively, after FDR correction, Figure 4E) in TT, as well as the up-regulation of the SDH complex, can indicate a strong mitochondrial dysfunction, which is well captured by both proteomic datasets in combination with the metabolomic data. Furthermore, both MTBE-SP3 and autoSP3 datasets agreed on a significant down-regulation of the abundance of OGDH in TT compared to NAT (MTBE-SP3: t-value = 4.06, p-value = 0.005, after FDR correction; autoSP3: t-value = 5.7, p-value < 0.0001, after FDR correction), an enzyme of the TCA cycle converting α-keto-glutarate to succinyl-CoA, upstream of the SDHA complex in the TCA cycle (Figure 4C), confirming a mitochondrial dysfunction. The integrated analysis of the proteomics and metabolomics datasets by ocEAn gives an additional perspective that is not directly recapitulated by a GO analysis of the proteomics dataset: The GO analysis mainly resulted in enriched terms related to RNA processing, gene expression, and translation (Supplementary Figure 5). In the GO analysis of the autoSP3 dataset, seven terms in the category ‘Biological Process’ were related to mitochondrial processes linked to mitochondrial gene expression or translation, but no terms were linked to mitochondrial metabolism. For the ocEAn results, both datasets also agreed on the up-regulation of the PKM enzyme in TT, which is the final rate-limiting step of glycolysis (MTBE-SP3: t-value = 5.76, p-value < 0.0001, after FDR correction; autoSP3: t-value = 4.63, p-value < 0.0001, after FDR correction). Finally, the ocEAn scores estimated from the metabolomic data showed slightly higher correlation coefficients with the proteomic data of the MTBE-SP3 dataset than the autoSP3 dataset (MTBE-SP3/ocEAn Pearson correlation: r = 0.45, p-value = 0.05; autoSP3/ocEAn Pearson correlation: r = 0.36, p-value = 0.12). Thus, despite some sparse differences between autoSP3 and MTBE-SP3, the two methods performed equally well, leading to the same biological insight in an integrated proteomic and metabolomic analysis of clinical samples (Figure 4A).

## Discussion

In general, the choice of extraction and processing method can highly influence downstream metabolomics and proteomics analysis of samples (Andresen *et al*, 2022). Depending on the composition and combination of solvents, the position of phase shifts, e.g., chloroform extraction (BLIGH & DYER, 1959) results in a lower phase containing lipids, an interphase containing proteins and an upper phase containing polar metabolites. Here, we applied a metabolite extraction suitable for broad metabolic profiling that also contains lipids by combining both polar and apolar phases. Following an adjusted biphasic extraction using 75% ethanol as organic solvent and MTBE as a substitute for chloroform, proteins will be precipitated as a pellet while the two resulting phases can be transferred, combined and dried for the metabolic profiling. We expect that a protein pellet instead of a protein interphase will produce a more discrete entity that can be collected to produce more consistent data in a downstream proteomic analysis. Similarly, an adjacent metabolite and lipid phase without an interfering protein-containing interphase can be handled more easily to produce more reliable results. Ultimately, this will allow to automate the metabolite extraction as no protein interphase is present. We previously showed that the usage of MTBE as an extraction buffer results in high-coverage, robust, and reproducible measurements of the metabolome compared to monophasic and other biphasic extractions (Gegner *et al*, 2022a). Besides the broad extraction range of polar metabolites and lipids, we here showed that the protein pellet obtained from the 75EtOH/MTBE extraction protocol can be readily integrated in already established down-stream processing steps (Müller *et al*, 2020) for proteome profiling.

To assess the performance of MTBE-SP3 workflow in comparison to autoSP3, we extracted bulk and/or cryo-pulverised and homogenised (powder) tissues and quantified their proteomes subsequently. The bulk samples come from physically distinct tissue pieces, while homogenised samples were taken from the same homogenate. We queried the proteomics datasets resulting from the two extraction methods (autoSP3, MTBE-SP3) and analysed the datasets to check for differences introduced by the preceding 75EtOH/MTBE extraction step. Both methods showed similarly low CV values for the different biological matrices (Table 1), indicating that MTBE-SP3 can be applied to a broad range of samples, and do not exhibit higher variability when measuring technical replicates. This result generally underlines the conclusion that autoSP3 and MTBE-SP3 quantify robustly the proteome of biological samples. We also scrutinised if MTBE-SP3 discriminates differently against physico-chemical properties of proteins looking at GRAVY and isoelectric point scores calculated from amino acid sequences. High similarity of physico-chemical properties indicated that MTBE-SP3 and autoSP3 exhibit very similar extraction characteristics for most of the sample types. For homogenised tissue types (fresh-frozen powder or FFPE powder), serum, plasma and cells MTBE-SP3 showed a low number of significantly abundant protein features, while this was slightly higher for bulk tissue types (bulk fresh-frozen tissue, bulk FFPE tissue, lung cancer). The underlying difference in the number of significantly abundant protein features between bulk and homogenised tissues is possibly caused by the variability in tissue sample content when probing from adjacent tissue neighbourhoods, given that bulk samples represent physically distinct tissue pieces, while homogenised samples were pooled, cryo-pulverised and taken from the same homogenate.

In the lung adenocarcinoma cohort, autoSP3 and MTBE-SP3 picked up equivalent differences between TT and NAT indicating that MTBE-SP3 assesses to a similar extent the proteome compared to the established autoSP3 method. The integration of proteomic and metabolomic data from NAT and TT using ocEAn, showed that both proteomic datasets are coherent with a tumour tissue displaying mitochondrial dysfunction, notably with deregulations of OGDH, SDH family enzymes and PKM. The SDH up-regulation in combination with the depletion of OGDH can well explain the depletion of succinate observed in tumours compared to healthy tissue, as illustrated by the joint up-regulation of both the abundance and ocEAn score of SDHA in TT *vs.* NAT. Furthermore, depletion of OGDH has been shown to lead to the stabilisation of HIF1A (Burr *et al*, 2016), which notably controls the expression of PKM. In the case of OGDH, only the protein abundance is down-regulated in TT *vs.* NAT, while the ocEAn score does not indicate any apparent global metabolic imbalance around OGDH. This can indicate that in the comparison between TT and NAT, OGDH is not acting as a strong metabolic bottleneck as the SDH complex. Thus, the integration of the metabolomic and proteomic datasets paint the picture of a mitochondrial dysfunction in tumour samples with an up-regulation of SDH enzymes and down-regulation of OGDH, leading to the depletion of succinate and up-regulation of the glycolysis metabolic pathway through the up-regulation of the PKM enzyme. This result was not recapitulated in the global interpretation of the proteomics data using GO analysis, which powerfully illustrates the complementarity of mono and multi-omics analyses. Finally, we showed that the ocEAn scores calculated from the metabolomic data had a better correlation with the differential expression analysis results of the proteomic data of the MTBE-SP3 dataset than the autoSP3. This can be explained by the fact that for MTBE-SP3 the proteome and metabolome measurements originate from the same sample, while they come from a different sample for autoSP3.

Taken together, we have devised a new single-sample workflow MTBE-SP3 by combining autoSP3 together with the 75EtOH/MTBE extraction workflow for proteomics and metabolomics sample processing, respectively. The MTBE-SP3 workflow enables the simultaneous processing of a single sample of all biological matrices for both metabolomic and proteomic analyses, thereby bypassing the problem of inter-sample variability and enabling more robust interpretation from the combined analysis of these modalities. As continuation of the autoSP3 workflow, the combined workflow is particularly relevant to perform multi-omics profiling of rare and limited sample amounts. We expect that robust single-sample workflows, such as MTBE-SP3, will advance the combined analysis of multi-omics experiments including proteomics and metabolomics.

## Materials and Methods

### Sample types

#### Tissue samples

All FFPE samples were collected from a biopsy punch of archival Ewing sarcoma xenografts derived from human Ewing sarcoma cell lines. Tumour purity and tissue integrity was assessed by a pathologist before sample processing. For the fresh-frozen samples, mouse liver tissue was used. Tissues were cut into small pieces, pooled by sample type, and aliquoted for further processing. One part was directly used for the autoSP3 workflow, while the second part was subjected to biphasic 75EtOH/MTBE extraction followed by autoSP3 (MTBE-SP3). The third part was cryo-pulverised and further processed (Powder-SP3).

#### Cell Culture

Human U2OS osteosarcoma cells were purchased from the American Type Culture Collection (ATCC) and tested for mycoplasma. Cells cultured in Dulbecco’s modified Eagle’s medium (DMEM) high glucose supplemented with 10% fetal bovine serum (FBS), 100 U/ml penicillin, and 100 µg/ml streptomycin at 37°C with 5% CO_2_. Cells were harvested using 0.05% Trypsin/EDTA and centrifuged at 400xg for 3 min. Cells were suspended and washed twice with 1x PBS, counted, and aliquoted into 10 Eppendorf tubes (1.6 million cells each). Next, cells were centrifuged for 5 minutes at 1000xg to remove the excess of PBS. Cell pellets were always kept on ice and subsequently stored at -20°C until further processing.

#### Plasma and serum

Plasma and serum samples were generated by pooling EDTA-plasma and serum samples acquired from the *Deutsches Rotes Kreuz Blutspendedienst*. These pooled blood samples were mixed at 4°C and aliquots of 100 µl generated. All aliquots were snap-frozen in liquid N_2_ and stored at -80°C until processing.

#### Lung adenocarcinoma cohort

Tissue samples were provided by the Lung Biobank Heidelberg, a member of the accredited Tissue Bank of the National Center for Tumor Diseases (NCT) Heidelberg, the Biomaterial Bank Heidelberg, and the Biobank platform of the German Center for Lung Research (DZL). The local ethics committees of the Medical Faculty Heidelberg (S-270/2001 (biobank vote) and S-699/2020 (study vote)) approved the use of specimens and data. All patients (cohort overview see Supplementary Table 1) included in the study signed an informed consent and the study was performed according to the principles set out in the WMA Declaration of Helsinki.

Tumour and matched distant (> 5 cm) tumour-free lung tissue samples from patients with non-small cell lung cancer (NSCLC), who underwent therapy-naive resection for primary lung cancer at Thoraxklinik at University Hospital Heidelberg, Germany were collected between 2016 and 2017. Tissues were snap-frozen within 30 minutes after resection and stored at -80°C until the time of analysis. All diagnoses were made according to the 2015 WHO classification for lung cancer by at least two experienced pathologists.

For further processing, cryosections (10-15 µm each) were prepared for each patient. The first and the last sections in each series were stained with hematoxylin and eosin (H&E) and were reviewed by an experienced lung pathologist to determine the proportions of viable tumour cells, stromal cells, normal lung cell cells, infiltrating lymphocytes and necrotic areas. Only samples with a viable tumour content of ≥ 50% were used for subsequent analyses.

**Supplementary Table 1.**
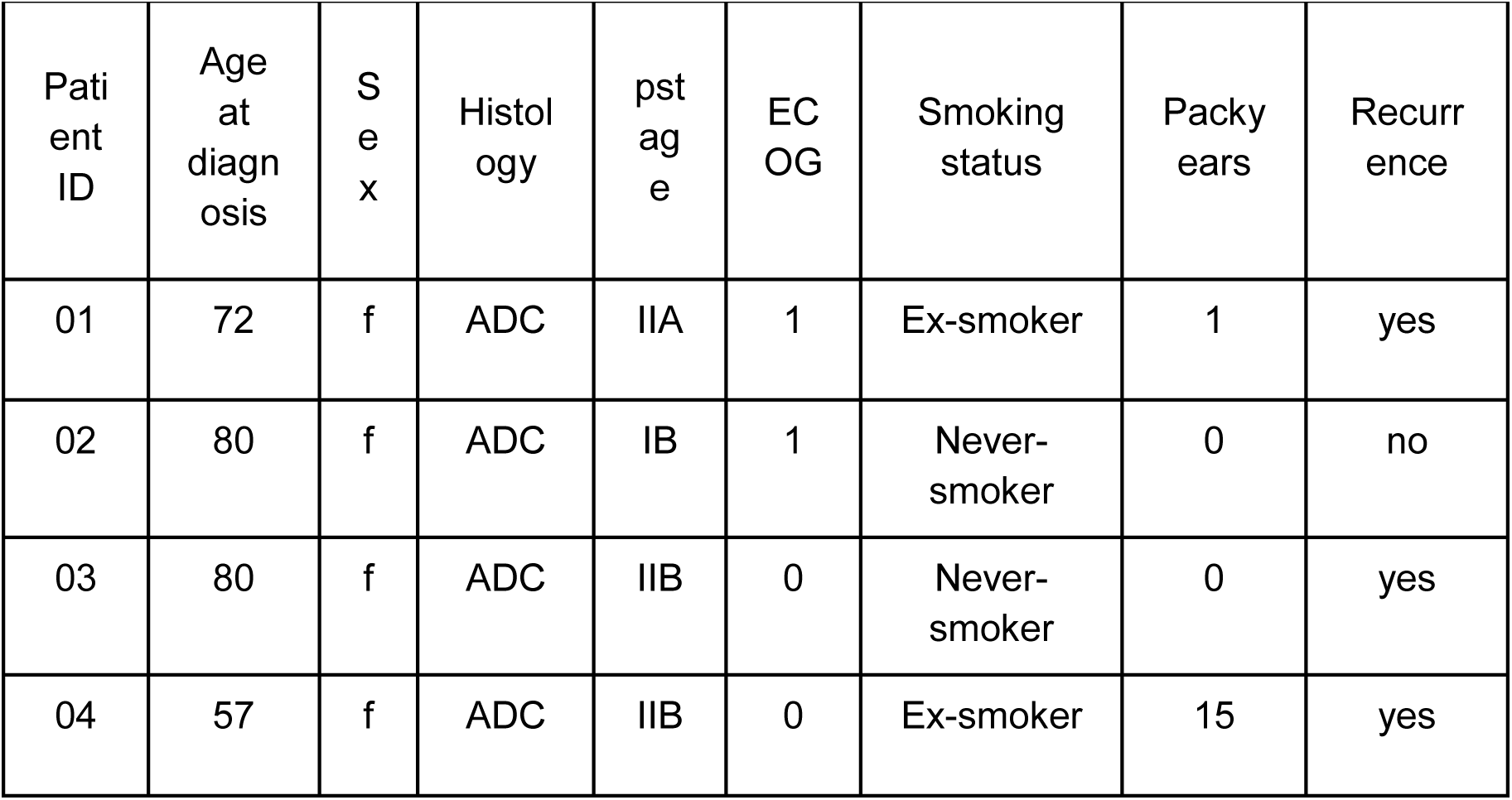
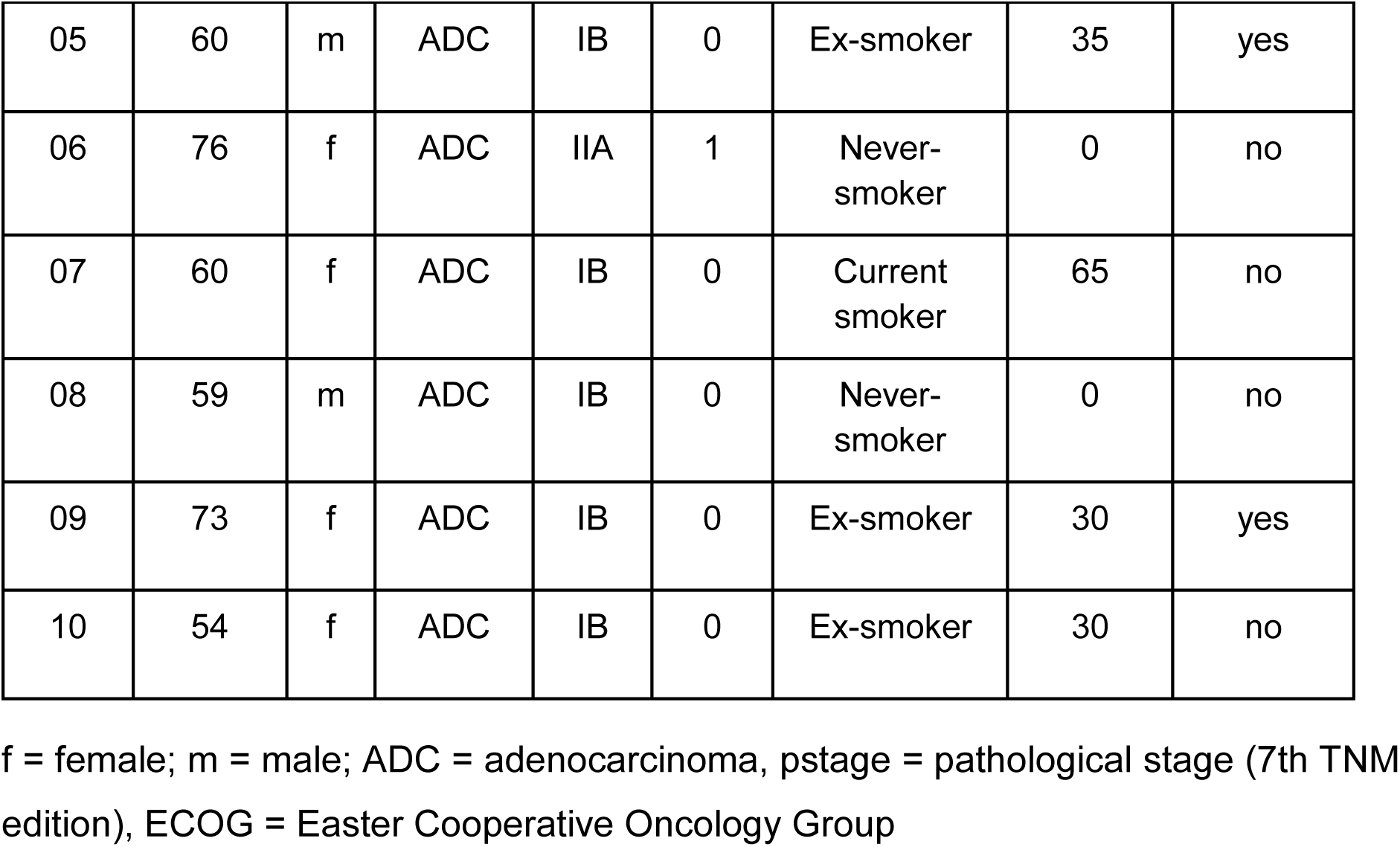
Information on lung adenocarcinoma patients.

#### Sample extraction

Tissue pieces were pulverised and extracted using an optimised protocol, specifically evaluated to produce broad coverage, high concentration and repeated values for tissue samples (Gegner *et al*, 2022a; Andresen *et al*, 2022). The biphasic 75EtOH/MTBE extraction generates two phases (containing polar metabolites and lipids) and additionally a protein pellet that was further analysed here (**Figure 1**). Briefly, samples were extracted using 300 µl ice-cold 75% ethanol, vortexed and sonicated for 5 min on ice or in the case of tissue, disrupted using a ball mill at 25 Hz for 30s. The resulting extract was mixed with 750 µl MTBE (tert-Butyl methyl ether) and kept at room temperature on a shaker (850 rpm) for 30 min. Next, 190 µl of H2O were added to separate the phases. The samples were vortexed and kept at 4°C for 10 min. Afterwards, the samples were centrifuged for 15 min at 13,000 g at 4°C. After the combination of both phases in the metabolite extraction, all samples were dried using an Eppendorf Concentrator Plus (at room temperature), stored at -80°C, and dissolved in 60µl isopropanol (30 µl of 100% isopropanol, followed by 30 µl of 30% isopropanol in water) before the measurement. The remaining protein pellet was kept at -80 °C until further processing using the autoSP3 proteomics workflow.

#### Standardised targeted metabolic profiling

Tissue extracts were processed following the manufacturer’s protocol of the MxP® Quant 500 kit (Biocrates). 10 µl of the samples or blanks were pipetted on the 96 well-plate-based kit containing calibrators and internal standards using an automated liquid handling station (epMotion 5075, Eppendorf) and subsequently dried under a nitrogen stream using a positive pressure manifold (Waters). Afterwards, 50 µl phenyl isothiocyanate 5% (PITC) was added to each well to derivatize amino acids and biogenic amines. After 1 h incubation time at room temperature, the plate was dried again. To resolve all extracted metabolites, 300 µl ammonium acetate (5 mM, in MeOH) were pipetted to each filter and incubated for 30 min. The extract was eluted into a new 96-well plate using positive pressure. For the LC-MS/MS analyses 150 µl of the extract was diluted with an equal volume of water. Similarly, for the FIA-MS/MS analyses 10 µl extract was diluted with 490 µl of FIA solvent (provided by Biocrates). After dilution, LC-MS/MS and FIA-MS/MS measurements were performed in positive and negative mode. For chromatographic separation an UPLC I-class PLUS (Waters) system was used coupled to a SCIEX QTRAP 6500+ mass spectrometry system in electrospray ionisation (ESI) mode. LC gradient composition and specific 50×2.1mm column are provided by Biocrates. Data was recorded using the Analyst (Version 1.7.2 Sciex) software suite and further processed via MetIDQ software (Oxygen-DB110-3005). All metabolites were identified using isotopically labelled internal standards and multiple reaction monitoring (MRM) using optimised MS conditions as provided by Biocrates. For quantification either a seven-point calibration curve or one-point calibration was used depending on the metabolite class.

#### Sample preparation for proteomic profiling

The sample preparation for proteome profiling was the same procedure for all sample types unless stated otherwise. A single cell suspension of U2OS cell aliquot was used as direct input into the standard method (Müller *et al*, 2020) or the biphasic MTBE/EtOH extraction. The latter resulted in a protein pellet which was resuspended in 1% SDS, 100 mM ammonium bicarbonate for further downstream processing using the autoSP3 method. Plasma and serum pools were aliquoted for the sample purpose to provide identical samples for both workflows, autoSP3 and MTBE-SP3. For fresh-frozen tissue, chunks were manually cut-off in the range of 1 to 3 mg as direct input into the standard autoSP3 method (Bulk-SP3). The remaining tissue (∼20-30 mg) was cryo pulverised and further aliquoted into equal proportions of powder. The powder was then either resuspended in 1% SDS, 100 mM ammonium bicarbonate and processed through autoSP3 (Powder-SP3) or subjected to the 75EtOH/MTBE extraction followed by autoSP3 (MTBE-SP3). Formalin-fixed and paraffin-embedded (FFPE) biopsy pillars (1 mm diameter and 8 mm length) were cut into cubes of roughly 1 mm^3^. Individual FFPE cubes were used as direct input into the standard autoSP3 method (Bulk-SP3) or a pool of cubes was used for cryo pulverisation. The resulting powder was aliquoted and resuspended in 1% SDS and 100 mM ammonium bicarbonate. The suspension was further processed through autoSP3 (Powder-SP3) or subjected to the 75EtOH/MTBE extraction followed by autoSP3 (MTBE-SP3). In summary, all sample types and formats (bulk, powder, or MTBE-pellet) were resuspended in 1% SDS and 100 mM ammonium bicarbonate and subjected to AFA-ultrasonication in a Covaris LE220plus instrument at the following settings: Duration 300 [seconds], PIP 450, DF 50, CPB 600, AIP 225 and dithering in Y +/- 1 mm, Z +/- 3 mm direction with 20 mm/second. Subsequently, the extracted amount of protein per sample was quantified using a BCA assay (Pierce) except for FFPE samples containing paraffin. FFPE samples were subjected twice to the sonication step interspaced by 2 cycles of heating at 95°C for 1 hour. Finally, all samples were processed through the autoSP3 protocol (Müller *et al*, 2020). For FFPE, additional wash steps (2x 200 µl 100% Isopropanol) and intermediate heating cycles of 10 minutes at 50°C were applied. Upon overnight proteolytic digestion, the resulting peptide samples were ready for injection into the mass spectrometer. Samples were stored at -20°C until measurement. The lung cancer fresh-frozen tissue cohort was processed via the bulk-SP3 and the (powder) MTBE-SP3 workflow.

#### Proteomic data acquisition

An equivalent of 200 ng peptides per sample were injected into a timsTOF Pro mass spectrometer (Bruker Daltonics) equipped with an Easy nLC 1200 system (Thermo) using the following method: peptides were separated using the Easy nLC 1200 system fitted with an analytical column (Aurora Series Emitter Column with CSI fitting, C18, 1.6 μm, 75 μm x 25 cm) (Ion Optics). The outlet of the analytical column with a captive spray fitting was directly coupled to a timsTOF Pro (Bruker) mass spectrometer using a captive spray source. Solvent A was ddH_2_O (Biosolve Chimie), 0.1% (v/v) FA (Biosolve Chimie), and solvent B was 80% ACN in dH_2_O, 0.1% (v/v) FA. The samples were loaded at a constant pressure. Peptides were eluted via the analytical column at a constant flow rate of 0.25 μL/min at 50°C followed by 10 minutes at 0.4 μL/min. During the elution, the percentage of solvent B was increased in a linear fashion from 4 to 17% in 15 min, then from 17 to 25% in 8 min, then from 25 to 35% in 5 min. Finally, the column was washed for 5 min at 100% solvent A. Peptides were introduced into the mass spectrometer via the standard Bruker captive spray source at default settings. The glass capillary was operated at 1600 V and 3 L/minute dry gas at 180°C. Full scan MS spectra with mass range m/z 100 to 1700 and a 1/k0 range from 0.85 to 1.3 V*s/cm2 with 100 ms ramp time were acquired with a rolling average switched on (10x). The duty cycle was locked at 100%, the ion polarity was set to positive, and the TIMS mode was enabled. The active exclusion window was set to 0.015 m/z, 1/k0 0.015 V*s/cm^2^. The isolation width was set to mass 700-800 m/z, width 2 – 3 m/z and the collision energy to 1/k0 0.85-1.3 V*s/ cm2, energy 27-45 eV. The resulting raw files were processed via MaxQuant (version 2.0.3.0) using the default settings unless otherwise stated. Label-free quantification (LFQ) and intensity-based absolute quantification (iBAQ) were applied using the default settings. Matching between runs was switched on.

#### Data processing for proteomics and metabolomics datasets

Data quality of protein and metabolite datasets was checked by MatrixQCvis (version 1.3.6, (Naake & Huber, 2022) and low-quality samples were excluded from further analysis. For the proteomics datasets (peptides for tissue comparison, proteins for tissue comparison, and proteins for lung adenocarcinoma cohort), LFQ intensities were log-transformed. The QC and PBS samples were excluded. For the lung cancer dataset, proteins with more than 18 from 35 measured values (i.e. no missing values) were retained in downstream analysis. For the metabolite dataset (lung adenocarcinoma cohort), the MetIDQ-derived dataset containing raw values was filtered according to the MetIDQ-derived quality scores such that metabolites that had at least ⅔ of valid values (i.e., 10x limit of detection and/or between the lower/upper limit of quantification).

#### Differential expression and overlap analysis for tissue dataset (peptides and proteins)

Differentially expression peptides and proteins were determined using limma (version 3.50.1) using lmFit (method = “ls”). Contrasts were specified via makeContrasts and fitted via contrasts.fit. The contrasts were defined as following: autoSP3 - MTBE-SP3 (cells), autoSP3 - MTBE-SP3 (Powder fresh-frozen tissue, contrast 1), autoSP3 - MTBE-SP3 (Bulk fresh-frozen tissue, contrast 2), autoSP3 - MTBE-SP3 (Powder FFPE tissue, contrast 1), autoSP3 - MTBE-SP3 (Bulk FFPE tissue, contrast 2), autoSP3 - MTBE-SP3 (plasma), and autoSP3 - MTBE-SP3 (serum). Moderated t-statistics of differential expression were determined by empirical Bayes moderation of the standard errors towards a global value using the eBayes function (using default values). Corresponding p-values were adjusted using FDR using the Benjamini-Hochberg method. α was set to 0.05.

The overlap between the different contrasts were analysed using functionality from the MatrixQCvis package (Naake & Huber, 2022) and visualised via functions from the upSetR package (Conway *et al*, 2017). Coefficient of variation (CV) was calculated via cv from MatrixQCvis (Naake & Huber, 2022) using the formula 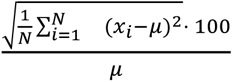 where μ is the mean of x.

#### Association of differential expressed peptides with physico-chemical properties for tissue dataset (peptides)

To calculate physico-chemical properties (isoelectric point and GRAVY scores of amino acids) we created the R package PhysicoChemicalPropertiesProtein that is available via https://github.com/tnaake/PhysicoChemicalPropertiesProtein. In brief, the ionizable groups of a protein/peptide sequence (N terminal, C terminal, δ-carboxyl group of glutamate, β-carboxyl group of aspartates, thiol group of cysteine, phenol group of tyrosine, imidazole side chains of histidine, ε-ammonium group of lysine, and guanidinium group of arginine) determine the isoelectric point of a given sequence. The pKA values are taken from (Kozlowski, 2016) and the implemented algorithm (bisection algorithm) is as in (Kozlowski, 2016). To calculate the isoelectric point the method IPC_protein was used. To calculate the GRAVY score, the hydropathy value for each residue is added and divided by the length of the sequence. The hydropathy values are taken from (Kyte & Doolittle, 1982). To test for association between physico-chemical properties and the extraction method (MTBE-SP3 *vs.* Bulk-SP3/Powder-SP3, autoSP3), Spearman’s rank correlation coefficient between GRAVY scores/isoelectric point and t-values from differential expression analysis of peptides were determined.

#### GO analysis for tissue dataset (proteins)

Protein ids were translated from UNIPROT to ENTREZ via AnnotationDbi (version 1.56.2). To this end, the following AnnotationDb objects were used: org.Hs.eg.db for cells, fresh-frozen tissue, FFPE tissue, plasma, and serum and org.Mm.eg.db for fresh-frozen tissue. Proteins that could not be translated to ENTREZ ids were removed from the downstream analysis. Over-representation of gene ontology (GO) terms was tested using the goana function from limma (version 3.50.1) where differential expressed proteins were proteins with adjusted p-values < 0.05 from differential expression analysis and the universe were all proteins present in the set.

#### Data analysis for adenocarcinoma lung cancer dataset (proteomics)

Protein IDs were translated from UNIPROT to SYMBOL via AnnotationDbi (version 1.56.2). Proteins with no corresponding SYMBOL IDs were removed from downstream analysis. To test for differential expression, a mixed linear model was created via limma (version 3.50.1) using duplicateCorrelation and lmFit. The blocking variable was set to individual. Contrasts were specified via makeContrasts and fitted via contrasts.fit. The contrasts were defined as follows: to test for differences between the autoSP3 and MTBE-SP3 method (TT_autoSP3 - NAT_autoSP3)/2 - (TT_MTBE-SP3) - NAT_MTBE-SP3)/2; to test for differences between the autoSP3 and MTBE-SP3 method in NAT NAT_autoSP3 - NAT_MTBE-SP3; to test for differences between the autoSP3 and MTBE-SP3 method in TT TT_autoSP3 - NAT_MTBE-SP3; to test for differences between TT and NAT in autoSP3 TT_autoSP3 - NAT_autoSP3; to test for differences between TT and NAT in MTBE-SP3 TT_MTBE-SP3 - NAT_MTBE-SP3; TT: tumour tissue, NAT: non-tumorous adjacent tissue. Moderated t-statistics of differential expression were determined by empirical Bayes moderation of the standard errors towards a global value using the eBayes function (using default values). Corresponding p-values were adjusted using FDR using the Benjamini-Hochberg (BH) method. α was set to 0.05. To test for tissue heterogeneity, the dataset was split into autoSP3 and MTBE-SP3 samples. For each subset, we randomly split the subsets into equal partitions. The blocking variable was set to tissue type (encoding information on NAT and TT origin). The contrast was defined as random_group1 - random_group2 to test for differences between the two random groups. Moderated t-statistics of differential expression and adjusted p-values were determined as described above.

#### GO analysis for adenocarcinoma lung cancer dataset (proteomics)

Protein ids were translated from SYMBOL to ENTREZ via AnnotationDbi (version 1.56.2). To this end, the org.Hs.eg.db AnnotationDb object was used. Proteins that could not be translated to ENTREZ ids were removed from the downstream analysis. Over-representation of gene ontology (GO) terms was tested using the goana function from limma (version 3.50.1) where differential expressed proteins were proteins with adjusted p-values < 0.05 from differential expression analysis and the universe were all proteins present in the set.

#### Data analysis for adenocarcinoma lung cancer dataset (metabolomics)

To test for differential expression, a mixed linear model was created via limma (version 3.50.1) using duplicateCorrelation and lmFit. The blocking variable was set to individual. The contrast was specified via makeContrasts and fitted via contrasts.fit. The contrast was set to TT - NAT to test for differences between tumour tissue (TT) and non-tumorous adjacent tissue (NAT). Moderated t-statistics of differential expression were determined by empirical Bayes moderation of the standard errors towards a global value using the eBayes function (using default values). Corresponding p-values were adjusted using FDR using the BH method. α was set to 0.05.

#### Integrated analysis of proteomic and metabolomic datasets for adenocarcinoma lung cancer dataset

In order to perform a pathway enrichment analysis with proteomic and metabolomic data, the first step was to connect metabolites to their corresponding enzymes, and embed the metabolites and enzymes in their respective pathways. A ready-to-use reaction network based on recon3D was extracted from the cosmosR package. As a pathway ontology, we used the cancer hallmark pathway collection from MSigDB (https://www.gsea-msigdb.org/gsea/msigdb). The identified metabolites of the metabolomic dataset were associated with their corresponding enzymes according to the reaction network. The hallmarks of the enzymes were transferred to the corresponding metabolite. This resulted in a hallmark pathway ontology containing both genes and metabolites annotated with their corresponding pathway hallmarks. The pathway enrichment analysis was performed with data from 4 patients, which had full overlap of metabolomic data and proteomic data generated with the autoSP3 and MTBE-SP3 pipelines. Using decoupleR, we ran pathway enrichment analyses with the run_wmean function of decoupleR, from which the norm_wmean enrichment score was extracted. The enrichment scores represent the number of standard deviations away from the mean of an empirical null distribution of scores for a given hallmark. The enrichment scores were calculated from the data presented in three different configurations: (1) from the proteomic data alone, (2) from the integrated metabolomic and proteomic dataset and (3) from the proteomic and metabolomic data separately, and subsequent averaging of the proteomic and metabolomic enrichment scores. This procedure was performed twice, once with the autoSP3 proteomic dataset, and once with the MTBE-SP3 proteomic dataset. For each dataset, the log2 fold change of protein and metabolic abundance were estimated individually for each of the 4 considered patients between the healthy and tumour samples. The fold changes of each protein and metabolite were then converted to z-scores across the 4 patients. Those z-scores were used as input for the decoupleR run_wmean function to estimate hallmark enrichment scores at the level of each patient. The enrichment scores obtained across the MSigDB hallmarks with the three data configurations were then correlated between the results of the autoSP3 and MTBE-SP3 datasets using Pearson correlation. All the scripts corresponding to this part of the analysis can be found at https://github.com/saezlab/prot_met_workflow/blob/main/scripts/create_combined_metab_gene_hallmarks.R, https://github.com/saezlab/prot_met_workflow/blob/main/scripts/SMARTCARE_decoupleR_sample_preparation.R and https://github.com/saezlab/prot_met_workflow/blob/main/scripts/SMARTCARE_decoupleR_pathway_enrichment_analysis.Rmd The ocEAn R package was used following the tutorial available at https://github.com/saezlab/ocean/blob/master/vignettes/tutorial_ocEAn.R The t-values of the metabolomic differential expression result (see Data analysis for adenocarcinoma lung cancer dataset (metabolomics)) were used as input for ocEAn. ocEAn distance penalty was set to 8, minimum branch length to 1 upstream and 1 downstream, and the ratio of upstream and downstream branch length for enzymes was left unbounded. The scores of reactions annotated as “reverse” were inverted. In order to compare the resulting metabolic imbalance scores of ocEAn with the proteomic data, multiple scores for the same enzyme (participating in different reactions) were averaged. For simplification purpose, we specifically restrained the interpretation of the results to enzymes of the canonical Kreb’s cycle (citrate -> isocitrate -> α-keto-glutarate -> succinyl-CoA -> succinate -> fumarate -> malate -> oxaloacetate -> citrate) with its incoming branch from glycolysis (phospho-enol pyruvate -> pyruvate -> acetyl-CoA) and its outgoing branch to acetyl-carnitine (acetyl-CoA + carnitine -> acetyl-carnitine). The averaged ocEAn metabolic imbalance score was then compared to the t-values from proteomic differential expression analysis (see Data analysis for adenocarcinoma lung cancer dataset (proteomics)), by computing the respective Pearson correlation coefficient of the averaged ocEAn scores with MTBE-SP3 and autoSP3 proteomic t-values, respectively. The script corresponding to this part of the analysis can be found here: https://github.com/saezlab/prot_met_workflow/blob/main/scripts/comparison_proteo <u>mic_ocEAn.R</u>

## Data and analysis script availability

The raw files, the search output files, as well as the utilised species databases have been deposited to the ProteomeXchange Consortium via the PRIDE partner repository under the following identifier: PXD046035

The analysis scripts are available via https://www.github.com/tnaake/MTBESP3_extraction_method

## Acknowledgements

We would like to thank Andrea Bopp and Martin Fallenbüchel from Lung Biobank Heidelberg as well as Felina Zahnow at the Division of Translational Pediatric Sarcoma Research (DKFZ) for their collection and processing of patient and xenograft specimens, respectively. We would like to thank David Robinson and Ben Warwick for providing the geom_flat_violin function. A.D., H.M.G., N.K.R., T.M., and T.N. and parts of the study were funded by the German Federal Ministry of Education and Research (BMBF) within the SMART-CARE consortium (fund no. 161L0212). The laboratory of T.G.P.G. is supported by the Barbara and Wilfried Mohr Foundation. The authors gratefully acknowledge the data storage service SDS@hd supported by the Ministry of Science, Research and the Arts Baden-Württemberg (MWK) and the German Research Foundation (DFG) through grant INST 35/1314-1 FUGG and INST 35/1503-1 FUGG.

## Supplementary Figures

**Supplementary Figure S1:**
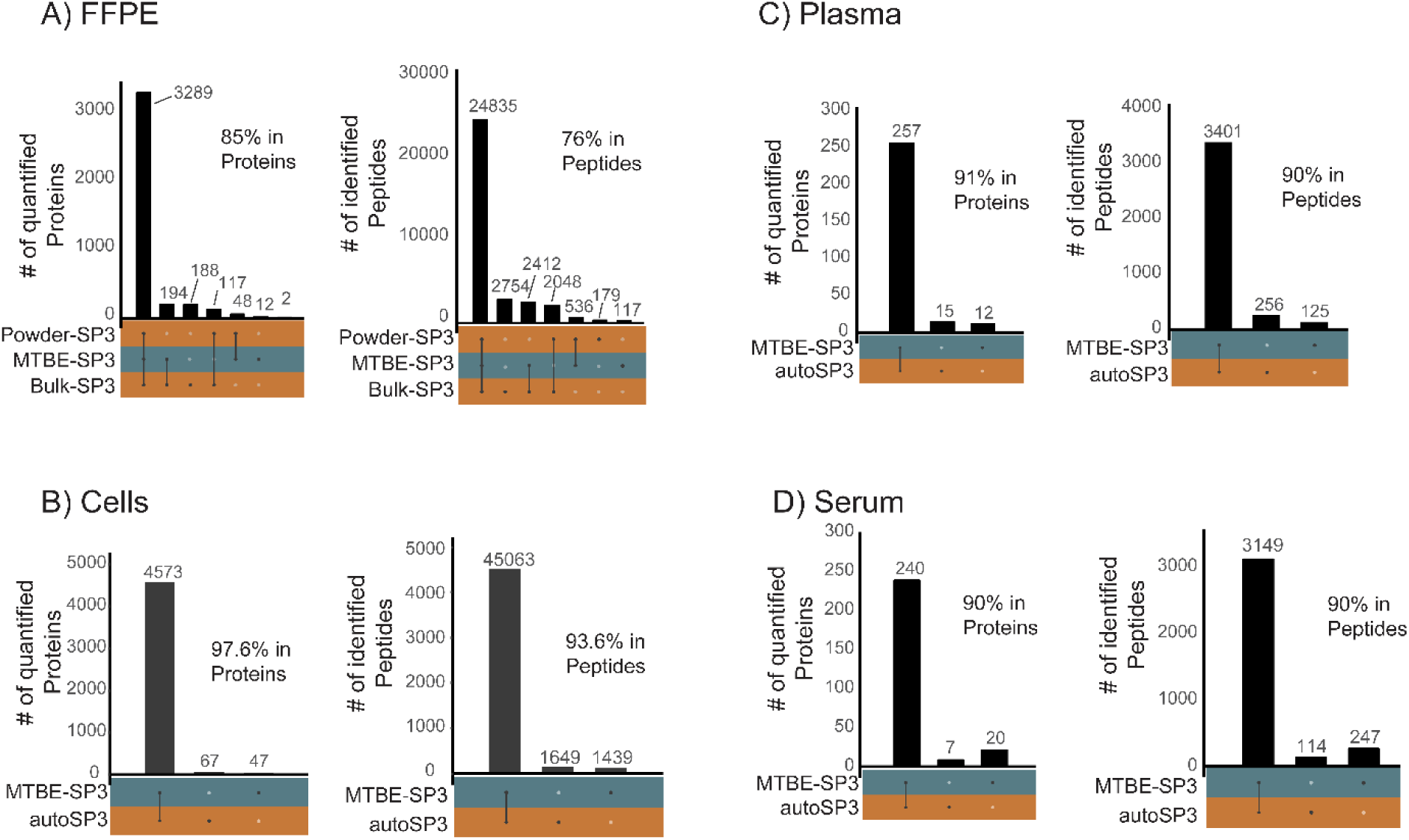
Overlap of extracted proteins and peptides in FFPE, cells, plasma, and serum samples. Joint and disjoint proteins and peptide sets. A) FFPE tissue. About 85% of proteins and 76% of peptides were detected in the joint set autoSP3 (Powder, Bulk) and MTBE-SP3. B) Cells. About 97.6% of proteins and 93.6% of peptides were detected in the joint set autoSP3 and MTBE-SP3. C) Plasma. About 91% of proteins and 90% of peptides were detected in the joint set autoSP3 and MTBE-SP3. D) Serum. About 90% of proteins and 90% of peptides were detected in the joint set autoSP3 and MTBE-SP3.

**Supplementary Figure S2:**
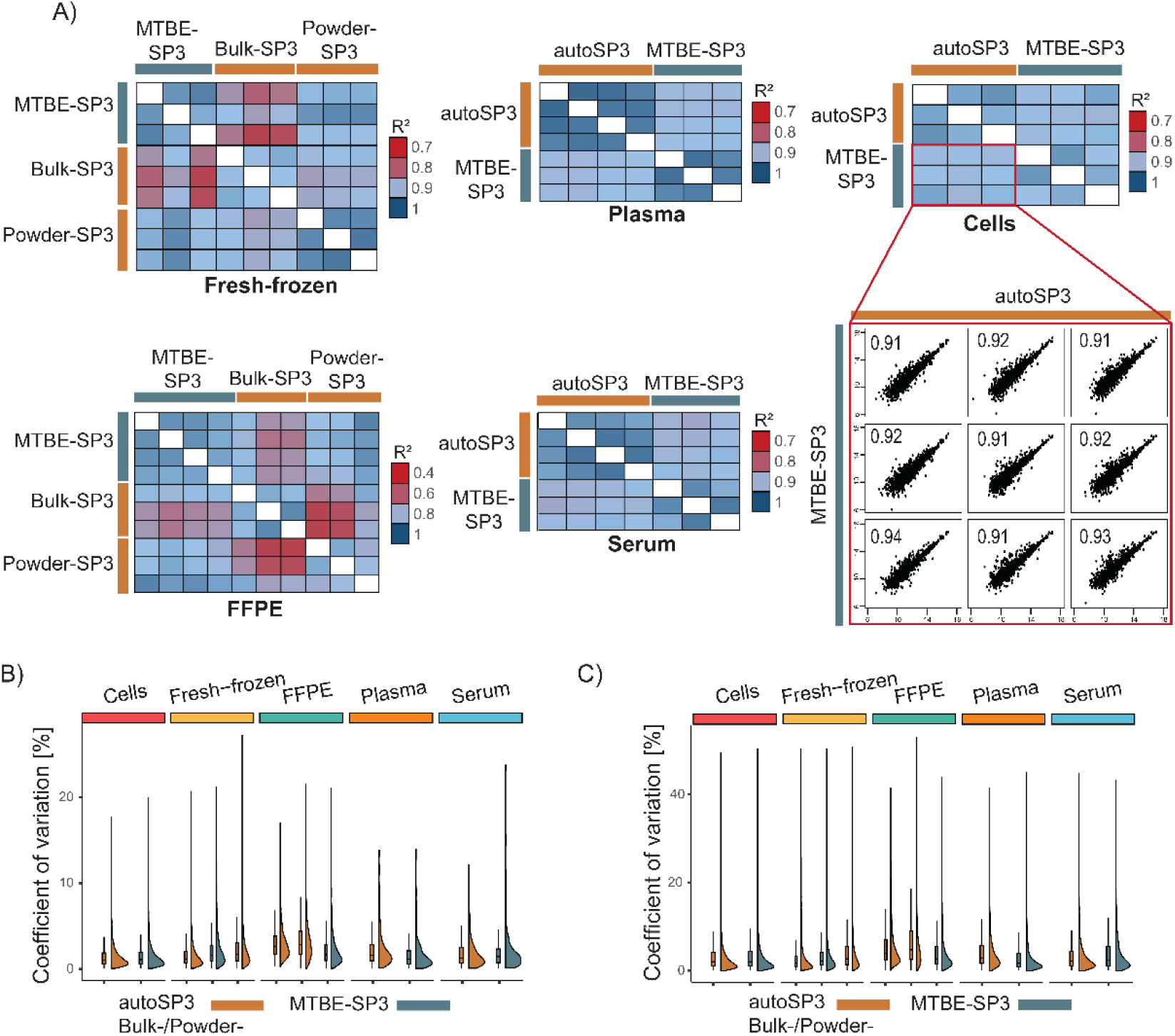
Comparison of autoSP3 and MTBE-SP3. A) Explained variance (R^2^) between log-transformed intensities of technical replicates. autoSP3 and MTBE-SP3 show overall high R^2^ between log-transformed intensities of replicates in all sample types as exemplified by the scatter plot for log-transformed intensities of autoSP3 and MTBE-SP3 in cells. The numerical values within the subpanel denotes the R^2^ between autoSP3 and MTBE-SP3 technical replicates. B) CV values of log-transformed protein intensities of technical replicates for autoSP3 and MTBE-SP3. MTBE-SP3 shows CV values in a similar range to autoSP3 in all sample types. C) CV values of log-transformed peptide intensities of technical replicates for autoSP3 and MTBE-SP3. MTBE-SP3 shows CV values in a similar range to autoSP3 in all sample types. CV: coefficient of variation.

**Supplementary Figure S3:**
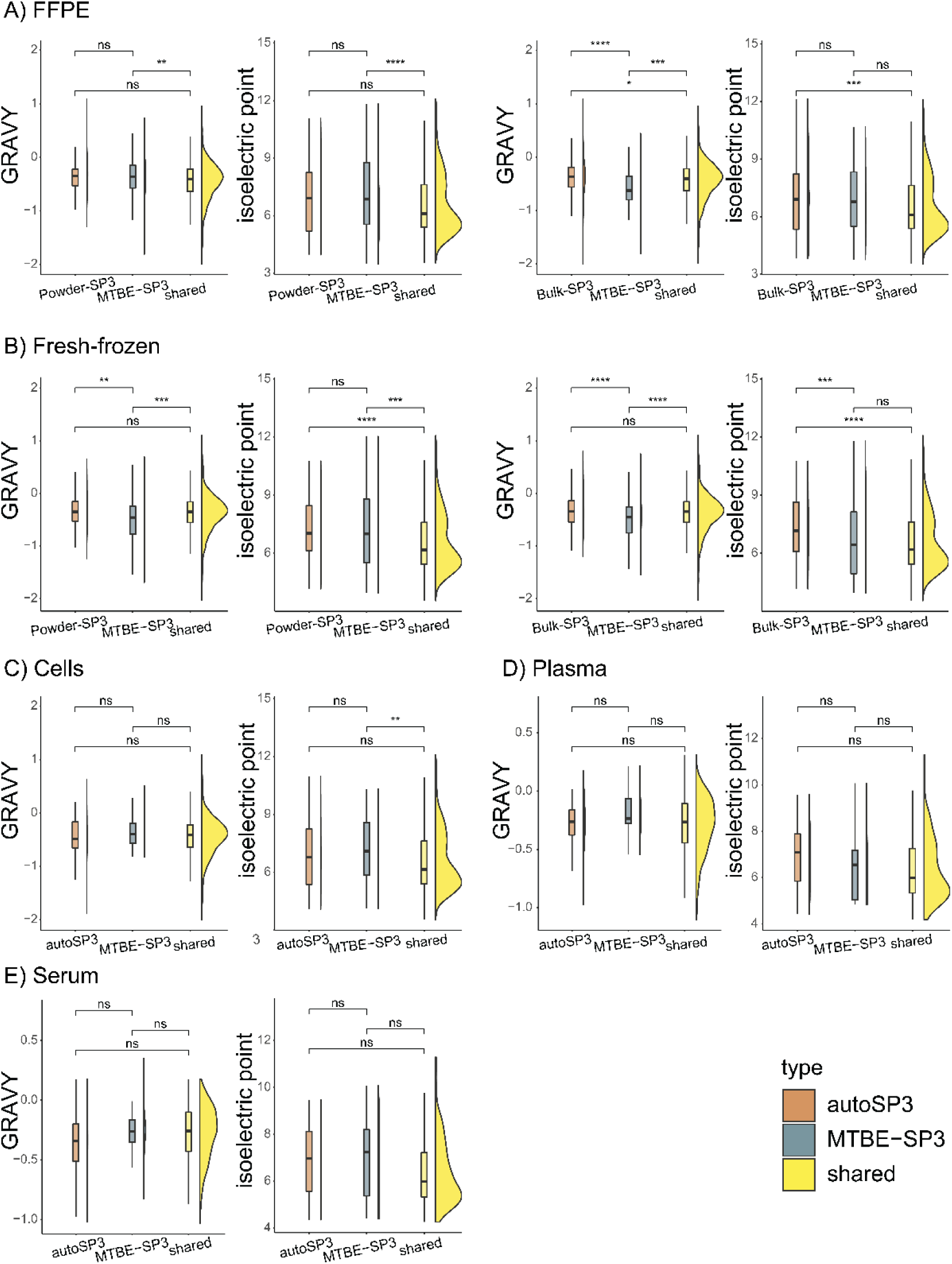
GRAVY and isoelectric point scores for proteins for the sets autoSP3/MTBE-SP3. Differences in means of GRAVY and isoelectric point values between the protein sets that were unique to autoSP3/MTBE-SP3 or shared (common) between autoSP3 and MTBE-SP3 were tested by the Wilcoxon signed-rank test (no adjustment for multiple testing). The effect size of differences was generally small, but statistically significant due to the high number of proteins. A) FFPE tissue. The two subfigures on the left refer to the contrast ‘Powder-SP3 *vs.* Powder-MTBE-SP3’. The two subfigures on the right refer to the contrast ‘Bulk-SP3 *vs.* Powder-MTBE-SP3’. B) Fresh-frozen tissue. The two subfigures on the left refer to the contrast ’Powder-SP3 *vs.* Powder-MTBE-SP3’. The two subfigures on the right refer to the contrast ‘Bulk-SP3 *vs.* Powder-MTBE-SP3’. C) Cells. D) Plasma. E) Serum. *: p-value < 0.05, **: p-value < 0.01, ***: p-value < 0.001, ****: p-value < 0.0001.

**Supplementary Figure S4:**
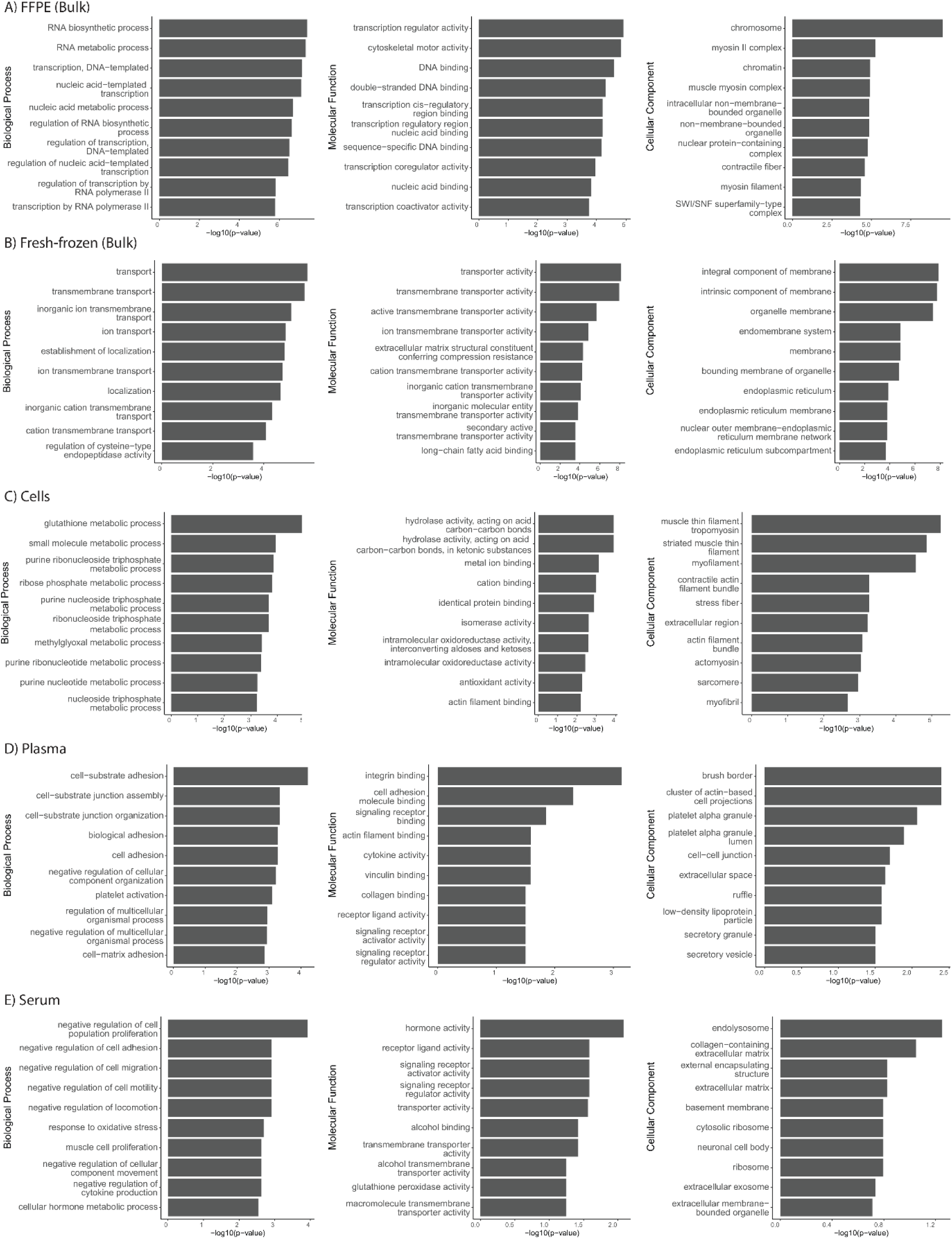
Enriched GO terms of differentially expressed proteins for FFPE (bulk, A) and fresh-frozen tissue (bulk, B), cells (C), plasma (D), and serum (E) between autoSP3 and MTBE-SP3 extraction. Shown are the top 10 terms for the categories Biological Process, Molecular Function, and Cellular Compartment.

**Supplementary Figure S5:**
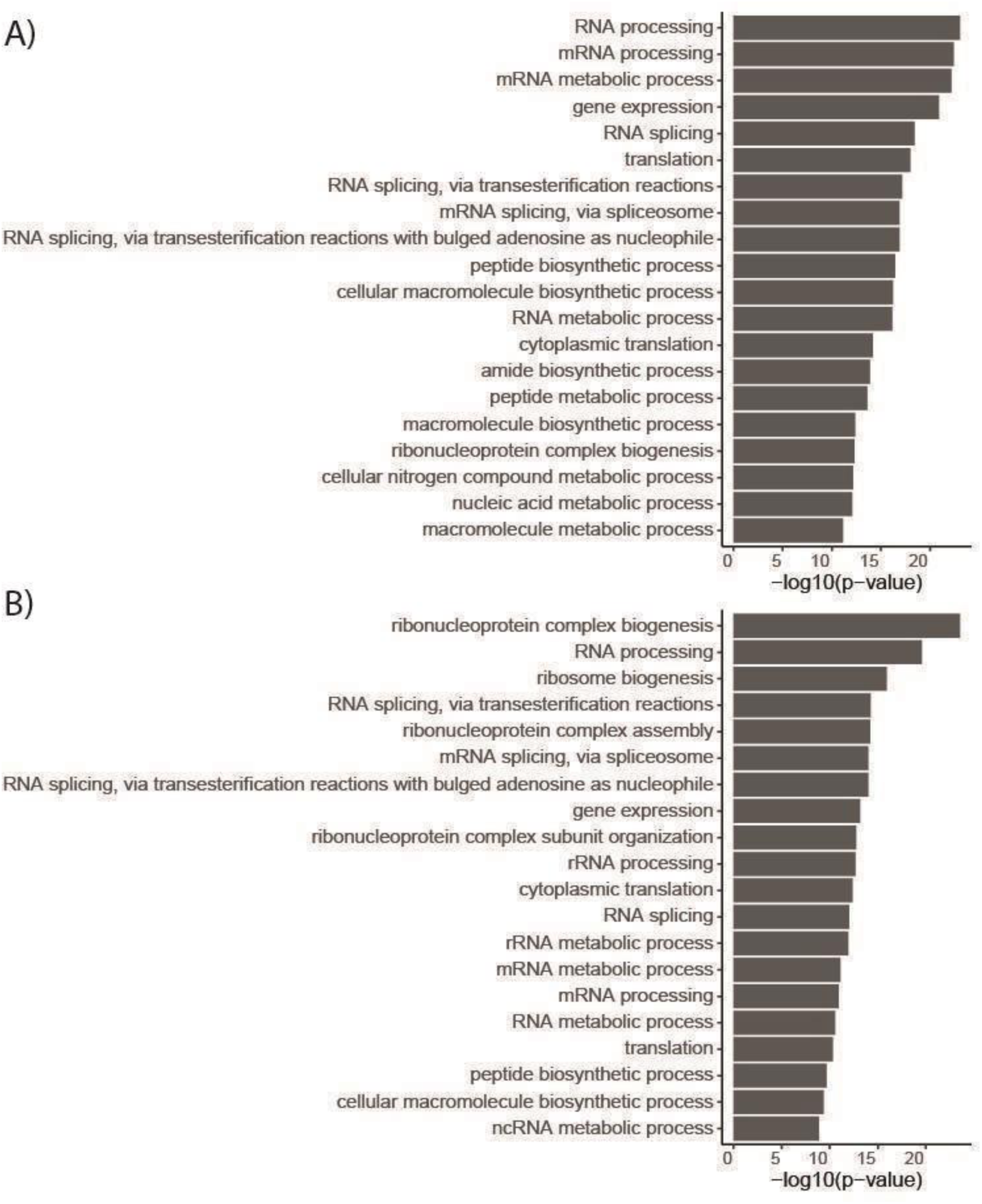
Enriched GO terms of differentially expressed proteins for the contrast TT *vs.* NAT in the lung adenocarcinoma dataset. A) GO terms for proteomics dataset acquired using the autoSP3 extraction. B) GO terms for proteomics dataset acquired using the MTBE-SP3 extraction. Shown are the top 20 terms for the category Biological Process. NAT: non-tumorous adjacent tissue. TT: tumorous tissue.

## List of Abbreviations

SP3: automated single-pot solid-phase-enhanced sample preparation
MTBE: Methyl-tert-butylether
LC-MS/MS: liquid chromatography-tandem mass spectrometry
FFPE: formalin-fixed paraffin-embedded
LFQ: label-free quantification
DDA: data-dependent acquisition
TT: tumorous tissue
NAT: non-tumorous adjacent tissue

